# Developmental Regulation of an Organelle Tether Coordinates Mitochondrial Remodeling in Meiosis

**DOI:** 10.1101/376509

**Authors:** Eric M. Sawyer, Pallavi R. Joshi, Luke E. Berchowitz, Elçin Ünal

## Abstract

Cellular differentiation involves remodeling cellular architecture to transform one cell type to another. By investigating mitochondrial dynamics during meiotic differentiation in budding yeast, we sought to understand how organelle morphogenesis is developmentally controlled in a system where regulators of differentiation as well as organelle architecture are known, but the interface between them remains unexplored. We found that mitochondria abruptly detach from the cell cortex shortly before segregating into gametes. Mitochondrial detachment is enabled by the programmed destruction of the **m**itochondria-**e**ndoplasmic reticulum-**c**ortex **a**nchor (MECA), an organelle tether that forms contact sites between mitochondria and the plasma membrane. MECA regulation is governed by a meiotic transcription factor, Ndt80, which promotes the activation of a conserved kinase, Ime2. We found that MECA undergoes Ime2-dependent phosphorylation. Furthermore, Ime2 promotes MECA degradation in a temporally controlled manner. Our study defines a key mechanism that coordinates mitochondrial morphogenesis with the landmark events of meiosis and demonstrates that cells can developmentally regulate tethering to induce organelle remodeling.

## INTRODUCTION

Mitochondria are essential organelles that host an array of cellular processes, ranging from ATP production to iron-sulfur cluster assembly. In many cell types, mitochondria are organized into a network of interconnected tubules that is dynamically remodeled by fusion and fission (Friedman and Nunnari, 2014). In addition, the position and motility of mitochondria are regulated to allow proper distribution within the cell and inheritance during cell division (Mishra and Chan, 2014; Westermann, 2014). Although the list of factors that modulate mitochondrial architecture and dynamics continues to expand, relatively little is known about their developmental regulation.

Fusion, fission, anchoring, and transport collectively shape the mitochondrial network. All of these processes are broadly conserved in eukaryotes but have been most extensively characterized in *Saccharomyces cerevisiae.* The budding yeast mitochondrial network exists as a branched structure that is dynamically remodeled by fusion and fission, while maintaining associations with the plasma membrane (Hoffmann and Avers, 1973; Nunnari et al., 1997). Plasma membrane anchoring requires a protein complex called MECA, for **m**itochondria-**E**R-**c**ortex **a**nchor (Cerveny et al., 2007; Klecker et al., 2013; Lackner et al., 2013; Ping et al., 2016). MECA belongs to a growing list of protein complexes collectively known as tethers, which establish membrane contact sites between disparate organelles (Elbaz-Alon et al., 2015; Elbaz-Alon et al., 2014; Kornmann et al., 2009; Murley and Nunnari, 2016; Murley et al., 2015). By physically bridging organelles, tethers enable interorganelle communication and establish spatial cellular organization (Eisenberg-Bord et al., 2016; Murley and Nunnari, 2016). Studies in multiple organisms have demonstrated the physiological importance of organelle tethers in controlling metabolism, intracellular signaling, pathogen defense, and organelle inheritance (Eisenberg-Bord et al., 2016; Helle et al., 2013; Prinz, 2014; Schrader et al., 2015). Furthermore, it has been shown that organelle tethers can be dynamically regulated in response to changes in cellular milieu, including metabolites and ions (Honscher et al., 2014; Kumagai et al., 2014; Nhek et al., 2010). However, whether and how these structures are subject to developmental regulation to meet the demands of differentiation into new cell types is not clear.

The principal cellular differentiation program in budding yeast is gametogenesis, which includes segregation of chromosomes by meiosis and the production of specialized gamete cells called spores. Various organelles, including mitochondria, undergo extensive remodeling during this process (Fuchs and Loidl, 2004; Gorsich and Shaw, 2004; Miyakawa et al., 1984; Neiman, 1998; Stevens, 1981; Suda et al., 2007; Tsai et al., 2014). Mitochondrial distribution changes dramatically during the meiotic divisions, when mitochondria lose their plasma membrane association, instead localizing near the gamete nuclei (Gorsich and Shaw, 2004; Miyakawa et al., 1984; Stevens, 1981). Subsequently, ~50% of the mitochondria from the progenitor cells is inherited by the gametes (Brewer and Fangman, 1980), and the residual pool is eliminated (Eastwood et al., 2012; Eastwood and Meneghini, 2015). Although little is understood about the mechanisms responsible for mitochondrial reorganization and inheritance during meiosis, many other aspects of this developmental program, including transcriptional and cell cycle control, have been worked out in great detail in this organism (Marston and Amon, 2004; Neiman, 2011; van Werven and Amon, 2011; Winter, 2012). To what extent the previously identified meiotic regulators control mitochondrial dynamics and segregation has been unexplored.

Here, we elucidated how mitochondrial reorganization is coordinated with meiotic development. We observed that mitochondria abruptly detach from the plasma membrane at the onset of anaphase II. To identify the mechanism responsible for regulating mitochondrial detachment, we examined a series of meiotic mutants with defects in meiotic progression. To our surprise, central meiotic regulators, such as the cyclin-dependent kinase CDK1/Cdc28 and the anaphase-promoting complex, were entirely dispensable for mitochondrial detachment. Instead, we found that the transcription factor Ndt80 and the meiosis-specific kinase Ime2 dictate the timing of mitochondrial detachment. Ndt80 controls mitochondrial detachment by inducing the expression of Ime2 and promoting its kinase activity (this study; Benjamin et al., 2003; Berchowitz et al., 2013). Ime2 phosphorylates both subunits of the MECA complex *in vitro.* Furthermore, Num1 undergoes Ime2-dependent phosphorylation *in vivo.* Finally, we show that Ime2 promotes MECA proteolysis, and this timely destruction of MECA drives mitochondrial detachment. Our results indicate that organelle tethering can be developmentally regulated to facilitate organelle remodeling, a feature of many cellular differentiation programs.

## RESULTS

### Mitochondria detach from the plasma membrane in meiosis

To characterize the morphology and dynamic behavior of mitochondria during meiotic differentiation, we monitored cells that simultaneously expressed fluorescent markers of mitochondria (Cit1-GFP) and the nucleus (Htb1-mCherry) using time-lapse microscopy. Prior to the nuclear divisions, the mitochondrial network retained its characteristic morphology, existing as an interconnected tubular structure anchored to the cell cortex. Consistent with previous reports (Gorsich and Shaw, 2004; Miyakawa et al., 1984; Stevens, 1981), we found that mitochondria dissociated from the cell cortex during meiosis II (Figure 1A; Movie S1). We term this phenomenon “mitochondrial detachment.” Our data indicate that mitochondrial detachment occurs coincident with anaphase II. At the time of mitochondrial detachment, 68% of cells had begun anaphase II (Figure 1A). By 10 min after mitochondrial detachment, 90% of cells had initiated anaphase II.

**Figure 1.**
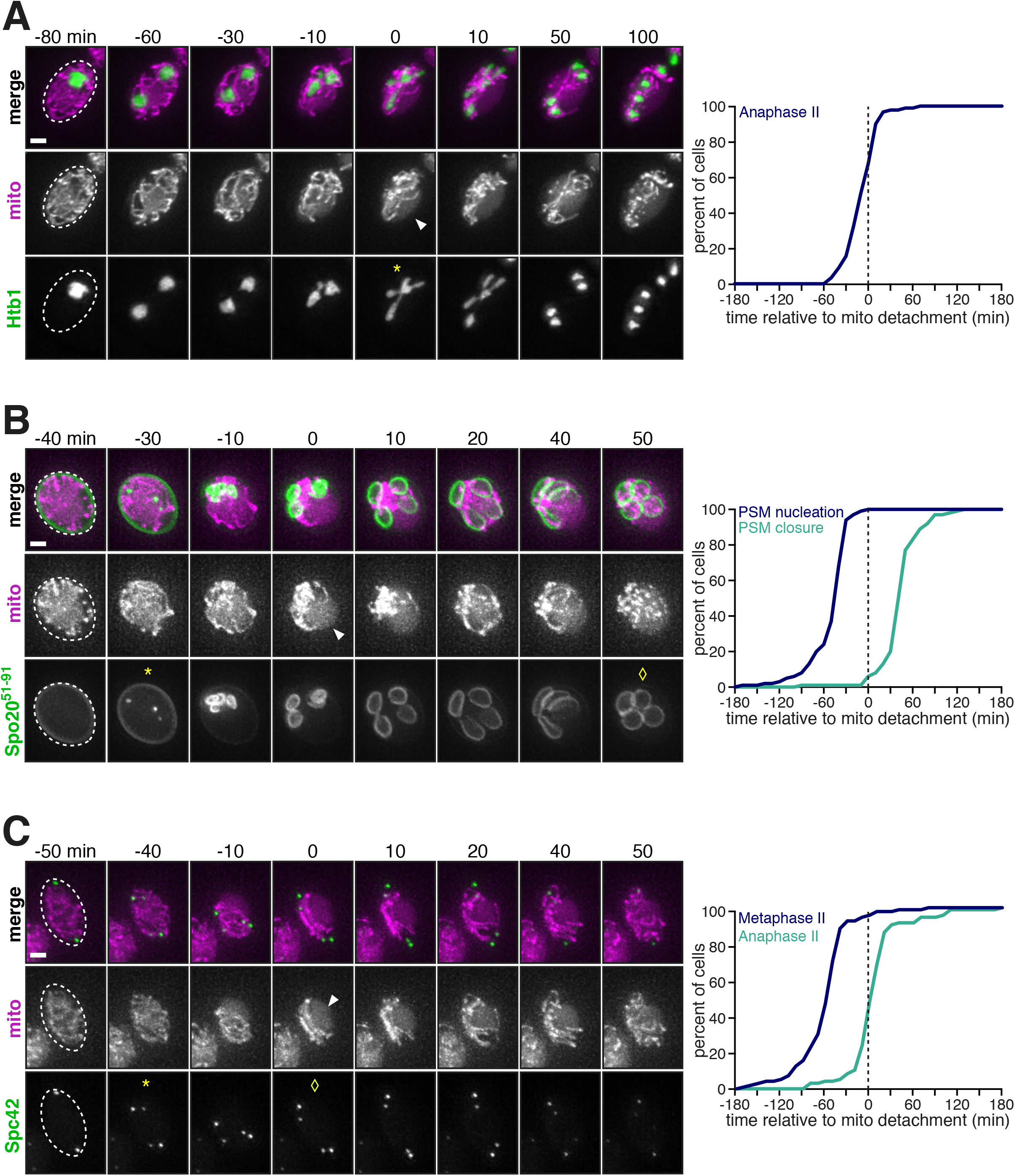
Mitochondria detach from the cell cortex during meiosis II. Movie montages and quantifications of cells expressing Cit1-GFP or Cit1-mCardinal to label mitochondria (mito) as well as a meiotic staging marker, imaged every 10 min. Mitochondrial detachment is defined as the loss of uniform, cortical distribution of mitochondria and indicated with an arrowhead. Cell boundaries are indicated with dashed lines. To determine the relative staging compared to other markers (below), mitochondrial detachment is defined to occur at 0 min. **A**. Mitochondrial detachment relative to the onset of the meiosis II nuclear division (anaphase II), marked by Htb1-mCherry (UB10257). Anaphase II is defined as the first appearance of a four-lobed nuclear morphology (*). *n =* 90. **B**. Mitochondrial detachment relative to prospore membrane nucleation and closure, marked by GFP-Spo20^51-91^ prospore membrane marker (UB13131). Prospore membrane nucleation is defined as the first appearance of Spo20^51-91^ puncta (*), and closure as the rounding up of fully elongated prospore membranes (◊). *n* = 100. **C**. Mitochondrial detachment relative to metaphase II and anaphase II, marked by Spc42-GFP (UB13129). Metaphase II is defined as the first appearance of two pairs of separated Spc42-GFP dots (*). Anaphase II is defined as the first appearance of concerted movement separating the sister spindle pole bodies in each pair (◊). *n* = 96. Scale bars, 2 *μ*m.

In order to further determine the timing of mitochondrial detachment, we used two additional staging markers. The first marker, GFP-Spo20^51-91^, is an indicator of plasma membrane biogenesis that takes place as part of gamete maturation (Nakanishi et al., 2004; Neiman, 2011). Concomitant with the meiosis I to meiosis II transition, this process, termed prospore membrane formation, begins with fusion of vesicles at the yeast centrosomes, known as spindle pole bodies. As judged by changes in GFP-Spo20^51-91^ localization, mitochondrial detachment occurred after membrane nucleation but prior to the closure of the newly formed plasma membranes (Figure 1B; Movie S2).

The second marker, Spc42-GFP, is a component of the spindle pole body. The distance between the duplicated spindle pole bodies is a reliable metric to determine the timing of metaphase to anaphase transition, since the spindle length increases approximately two-fold during this period (Kahana et al., 1995; Palmer et al., 1989; Yeh et al., 1995). We measured when mitochondrial detachment took place with respect to changes in spindle length in cells carrying Spc42-GFP and Cit1-mCardinal. This analysis revealed that mitochondrial detachment occurred at the beginning of anaphase II (Figure 1C; Movie S3). Hence, the timing of mitochondrial detachment is precise and occurs with stereotyped timing relative to other well-defined meiotic events.

### Many canonical cell cycle regulators are dispensable for mitochondrial detachment

Because mitochondrial detachment occurred simultaneously with anaphase II onset, we reasoned that cell cycle regulators with characterized meiotic functions might jointly control the meiotic divisions and mitochondrial detachment. Since the initial steps of spore formation occur during meiosis II, active coupling of chromosome and organelle segregation could ensure gamete fitness. We monitored mitochondrial detachment and meiotic progression in strains carrying deletion or conditional alleles of genes encoding key cell cycle regulators (Figure 2A). We also noted that at meiotic entry, all of the mutants examined showed mitochondrial morphology indistinguishable from wild type, indicating that these alleles did not constitutively alter mitochondrial organization (Figure 2B-H). 8 hours after induction of meiosis, the vast majority of wild-type cells contained four distinct nuclei that had not yet assembled into spores. In these cells, mitochondria invariably detached from the cortex and instead localized near the four post-meiotic nuclei (Figure 2B).

**Figure 2.**
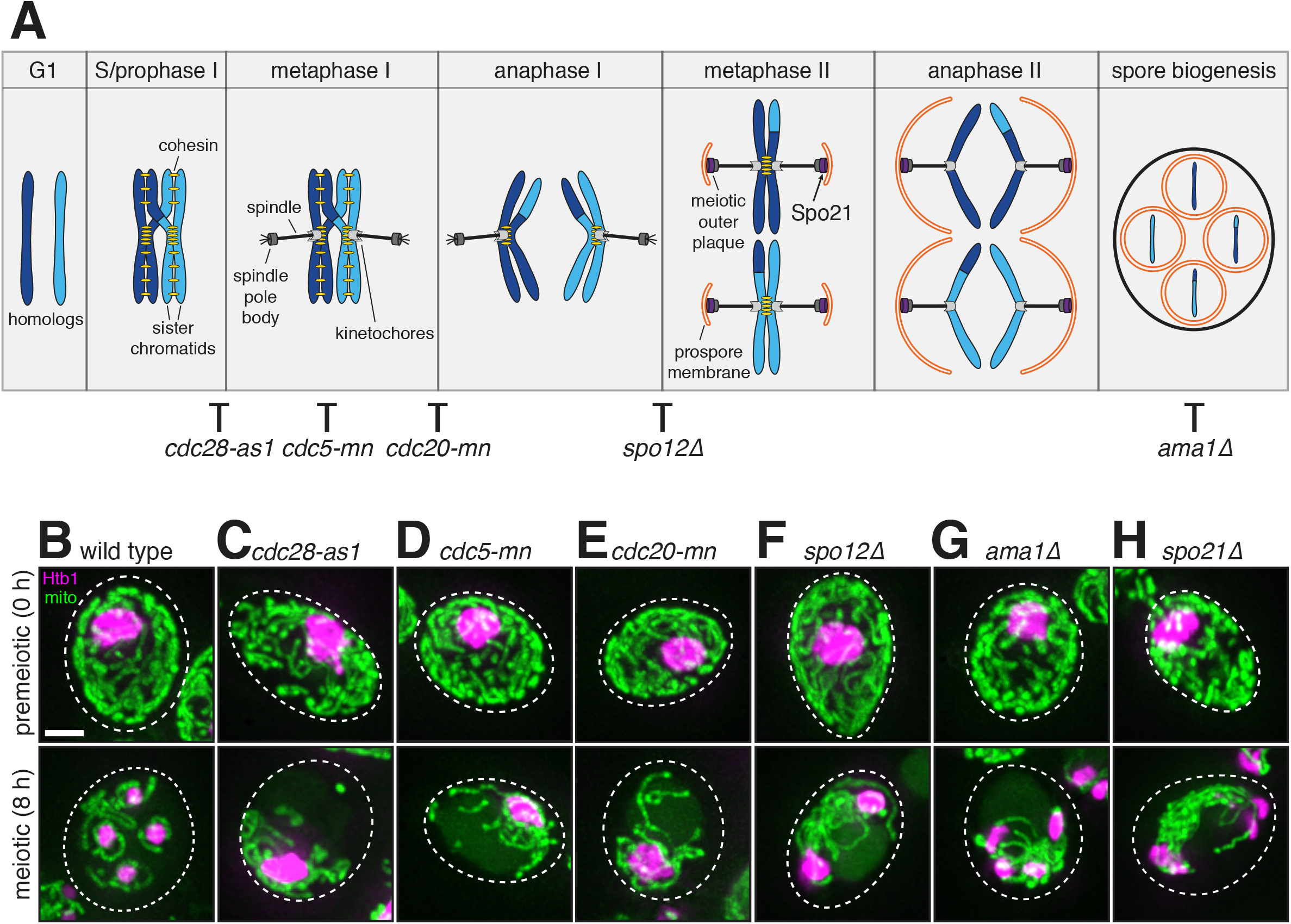
Mitochondrial detachment is uncoupled from the meiotic divisions and spore development. **A**. Schematic of meiotic chromosome segregation and spore development. Meiotic regulators and spore development genes are labeled at key stages for their functions, and where disruption of their function perturbs meiotic progression. **B-H**. Maximum intensity projections of fixed wild-type and mutant cells at 0 h in SPO (top) and 8 h in SPO (bottom). Mitochondria (mito), mitoGFP or Cit1-GFP. Nuclei, Htb1-mCherry. Cell boundaries are indicated with dashed lines. **B**. wild type (UB7155) **C**. *cdc28-as1* synchronized by *pGAL-NDT80 GAL4.ER* (UB9494), with 1 *μ*M 1-NM-PP1 and 1 *μ*M β-estradiol added simultaneously at 5 h **D**. *cdc5-mn,* which is *pCLB2-CDC5* (UB7278) **E**. *cdc20-mn,* which is *pCLB2-CDC20* (UB7343) **F**. *spo12Δ* (UB7345) **G**. *ama1Δ* (UB7533) H. *spo21Δ* synchronized by *pGAL-NDT80 GAL4.ER* (UB9239). Scale bar, 2 *μ*m.

Among the cell cycle regulators that we analyzed, the polo kinase Cdc5, the anaphase-promoting complex (APC) activator Cdc20, and the cyclin-dependent kinase Cdc28/Cdk1 are all essential for cell viability. To avoid perturbing the mitotic functions of these genes, we depleted *CDC5* and *CDC20* only from meiotic cells by replacing their promoters with the mitosis-specific *CLB2* promoter (Lee and Amon, 2003). To downregulate *CDC28* we utilized a chemical inhibitor sensitive allele, *cdc28-as1* (Bishop et al., 2000). It has been previously reported that each mutant perturbs meiotic chromosome segregation: *cdc5* and *cdc20* mutants are defective in exiting metaphase I (Lee and Amon, 2003), while inactivation of *cdc28-as1* in prophase I with 1-NM-PP1 inhibits meiosis I spindle assembly entirely (Benjamin et al., 2003). In each condition, the expected nuclear division defect was observed. However, mitochondrial detachment was unaffected. Mitochondria not only detached from the plasma membrane but also adopted their perinuclear localization, similar to wild-type cells (Figure 2C-E).

In addition to testing essential cell cycle regulators, we also assessed the role of nonessential regulators with defined meiotic functions. The Cdc14 Early Anaphase Release (FEAR) network controls the release of the Cdc28 antagonist phosphatase Cdc14 (Stegmeier et al., 2002; Yoshida et al., 2002). FEAR network signaling is absent in *spo12Δ* cells, which results in aberrant meiosis I spindle disassembly, culminating in the formation of binucleate post-meiotic cells instead of tetranucleate (Kamieniecki et al., 2005; Klapholz and Esposito, 1980; Marston et al., 2003). In *spo12Δ* cells, mitochondrial detachment was normal (Figure 2F). Mitochondrial detachment was also unaffected in *ama1Δ* cells (Figure 2G), which lack a meiosis-specific APC activator required for spore biogenesis (Cooper et al., 2000; Diamond et al., 2009).

Finally, we sought to address the possibility that prospore membrane formation is required for mitochondrial detachment, such as through sequestration of mitochondria into spores, since close proximity between mitochondria and prospore membrane has been previously observed (Suda et al., 2007). Synthesis of the prospore membrane requires assembly of a meiosis-specific structure on the cytoplasmic face of the spindle pole body, called the meiotic outer plaque (Knop and Strasser, 2000). In the absence of Spo21 (also known as Mpc70), a meiotic outer plaque component, other subunits fail to localize to the meiotic outer plaque, and therefore prospore membrane formation is completely disrupted (Knop and Strasser, 2000). We found that in *spo21Δ* cells, mitochondrial detachment was entirely unimpeded (Figure 2H). From these analyses, we conclude that much of the regulatory scheme that defines meiotic chromosome segregation and cellular differentiation are dispensable for mitochondrial detachment and that other factors must be involved in regulating when and how mitochondria dissociate from the cell cortex.

### The meiosis-specific transcription factor Ndt80 is required for mitochondrial detachment

We noted that in wild-type cells, mitochondrial morphology was indistinguishable at meiotic entry and prophase I, with mitochondrial detachment occurring abruptly during the second meiotic division. The master regulator controlling the transition to the meiotic divisions is the transcription factor Ndt80 (Chu and Herskowitz, 1998; Xu et al., 1995). Transcription of *NDT80* mRNA occurs during prophase I, but the ability of Ndt80 protein to localize to the nucleus is restricted by the pachytene checkpoint, which monitors the completion of double-strand break repair requisite for successful chromosome segregation (Chu and Herskowitz, 1998; Hepworth et al., 1998; Tung et al., 2000; Wang et al., 2011). In the absence of *NDT80,* cells exhibit a prolonged arrest during the pachytene stage of prophase I, failing to undergo meiotic divisions and subsequent gamete maturation (Xu et al., 1995).

To determine whether *NDT80* is required for mitochondrial detachment, we examined mitochondrial morphology in a *pGAL-NDT80* strain (Benjamin et al., 2003; Carlile and Amon, 2008). The *pGAL-NDT80* allele allows controlled induction of *NDT80* transcription by a β-estradiol activatable Gal4 fusion to the estrogen receptor protein (Gal4.ER) (Benjamin et al., 2003; Carlile and Amon, 2008). When we released cells from a 5-h Ndt80 block by addition of β-estradiol, mitochondrial detachment occurred normally (Figure 3A, +Ndt80). However, when the inducer was withheld, mitochondria remained persistently localized to the cell cortex (Figure 3A, -Ndt80), indicating that *NDT80* expression is necessary for mitochondrial detachment. *pGAL-NDT80* cells monitored by time-lapse microscopy showed identical behavior (Figure S1A; Movie S4 and S5). These experiments also confirmed that mitochondrial detachment is developmentally regulated and not an indirect outcome of prolonged nutritional deprivation. We conclude that Ndt80, a key regulator of meiotic events, is required for mitochondrial detachment.

**Figure 3.**
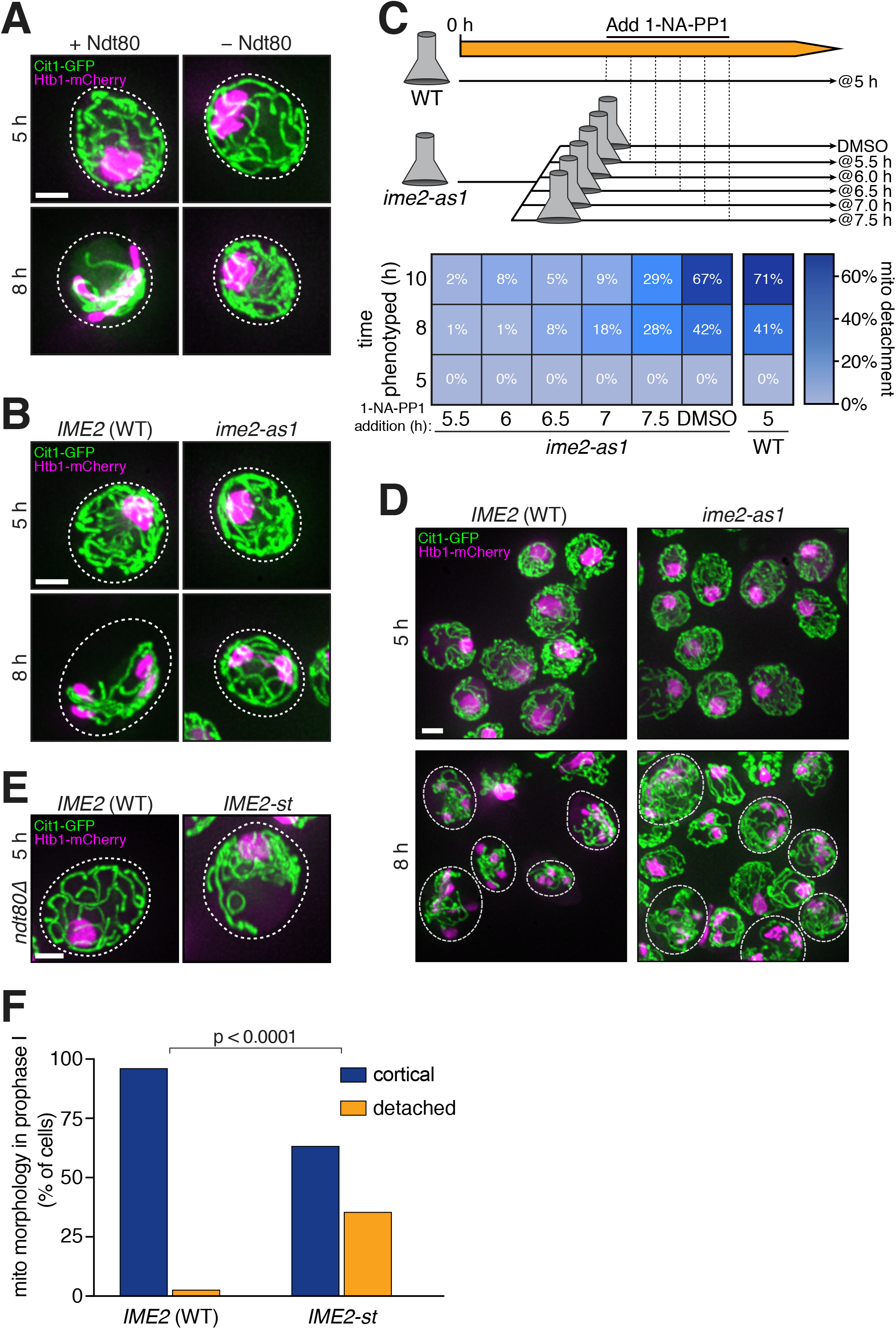
Ndt80 and Ime2 regulate mitochondrial detachment. **A**. Maximum intensity projections of fixed *pGAL-NDT80 GAL4.ER* cells (UB9496) with *NDT80* induced by addition of 1 *μ*M β-estradiol (+Ndt80) or ethanol vehicle control (-Ndt80). Sporulation cultures were arrested for 5 h, then split and subjected to the indicated treatment. Mitochondria, Cit1-GFP. Nuclei, Htb1-mCherry. Cell boundaries are indicated with dashed lines. **B**. Maximum intensity projections of fixed *pGAL-NDT80 GAL4.ER* cells containing wild-type (WT) *IME2* (UB9158) or *ime2-as1* (UB9844). Sporulation cultures were arrested for 5 h, then 1 *μ*M β-estradiol and 20 *μ*M 1-NA-PP1 were added simultaneously. Cell boundaries are indicated with dashed lines. **C-D**. Execution point time course, with the timing of Ime2 inhibition varied and *pGAL-NDT80* induced at 5 h. **C**. Top: Schematic of the execution point time course. Wild-type *pGAL-NDT80 GAL4.ER* (UB9158) and *pGAL-NDT80 GAL4.ER ime2-as1* (UB9844) were induced to sporulate and arrested for 5 h, then treated with 1 *μ*M β-estradiol to induce *NDT80.* Next, the *ime2-as1* culture was split and treated with 20 *μ*M 1-NA-PP1 at the indicated time points. DMSO served as a vehicle control and was added at 5 h. Bottom: Heat map showing the frequency of mitochondrial detachment at 5, 8, and 10 h after transfer to SPO medium (i.e., the time of fixation) for each of the execution points. *n* = 2200 total cells counted, with *n ≥* 32 per panel in the heat map. D. Maximum intensity projections from wild-type control and the 7.5 h execution point. Cell boundaries are indicated with dashed lines for meiosis II cells only. **E**. Maximum intensity projections of prophase I arrested cells (*ndt80Δ,* 5 h in SPO) containing wild-type *IME2* (UB17328) or *IME2-st* (UB17330). Cell boundaries are indicated with dashed lines. F. Quantification of mitochondrial morphology (cortical or detached) for the experiment in panel **E**. Fisher’s exact test p < 0.0001 (n = 100 per genotype). Scale bars, 2 *μ*m.

### Ime2 kinase is required for mitochondrial detachment

Ndt80 directly regulates the expression of approximately 200 genes (Cheng et al., 2018; Chu and Herskowitz, 1998). Among them, a particularly compelling candidate is the meiosis-specific kinase Ime2 (Foiani et al., 1996; Kominami et al., 1993; Nocedal et al., 2017; Smith and Mitchell, 1989; Yoshida et al., 1990). Ime2 belongs to a family of serine/threonine protein kinases displaying sequence similarities to both cyclin-dependent kinases and mitogen-activated protein kinases (Irniger, 2011). Although originally characterized as an early gene required for pre-meiotic S phase (Dirick et al., 1998), Ime2 kinase activity is significantly higher during meiosis II than any other meiotic stages (Benjamin et al., 2003; Berchowitz et al., 2013). It has been shown that Ndt80 is additionally required for the increase in Ime2 activity during meiosis II, independent of its role in regulating *IME2* expression (Benjamin et al., 2003; Berchowitz et al., 2013).

To determine whether Ime2 is necessary for mitochondrial detachment, we employed a conditional allele, *ime2-as1,* that can be selectively inhibited by the drug 1-NA-PP1 (Benjamin et al., 2003). By controlling the timing of inhibitor treatment, we could bypass the requirement for Ime2 in pre-meiotic S phase. In cells arrested in prophase I by the Ndt80 block, simultaneous addition of β-estradiol and 1-NA-PP1 resulted in retention of mitochondria at the cortex, even though cells performed meiosis I, as previously reported (Benjamin et al., 2003) (Figure 3B). Mitochondrial detachment was normal in *IME2* wild-type cells that were identically treated, ruling out non-specific effects of the drug treatment (Figure 3B). We obtained identical results by time-lapse microscopy (Figure S1B; Movie S6 and S7). These findings show that *IME2* function is necessary for mitochondrial detachment.

### Ime2 regulates mitochondrial detachment independent of its role in Ndt80 activation

Our experiments thus far indicate that Ndt80, a transcription factor, and Ime2, a kinase, are involved in mitochondrial detachment. Previous studies found that Ndt80 is required for the elevated activity of Ime2 during the meiotic divisions (Benjamin et al., 2003; Berchowitz et al., 2013), and conversely, Ime2 is required for the accumulation of Ndt80 and its full activation via phosphorylation (Benjamin et al., 2003; Sopko et al., 2002). Consequently, it was unclear whether the contribution of Ndt80 to the mitochondrial detachment phenotype was primarily by induction of Ime2, or the reverse model where the contribution of Ime2 is through enhancing Ndt80 activity. To distinguish between these possibilities, we first varied the timing of Ime2-as1 inhibition from 5.5 h to 7.5 h, keeping constant the time of *pGAL-NDT80* induction at 5 h. We found that mitochondrial detachment was acutely sensitive to Ime2 inactivation. The frequency of mitochondrial detachment showed a graded response over 30-min intervals of drug addition timing, with no drug addition time point recapitulating the high frequency of mitochondrial detachment observed in wild-type cells (Figure 3C). As reference, by mRNA-seq (Brar et al., 2012), Ndt80 target genes are induced within ~1 h of *pGAL-NDT80* induction in wild-type cells. Addition of 1-NA-PP1 at 7.5 h resulted in a mild effect on the meiotic divisions, yet mitochondrial detachment was still defective in a substantial fraction of the cells (Figure 3C and 3D). This result suggests that the contribution of Ndt80 to mitochondrial detachment is principally through the regulation of Ime2 activity.

Next, we tested whether Ime2 could promote mitochondrial detachment in cells lacking a functional *NDT80* gene (*ndt80Δ).* For this, we used an *IME2* allele that has elevated activity throughout meiosis (*IME2-st,* Sia and Mitchell, 1995). Even though the absence of *NDT80* expression completely blocked meiotic progression, ~40% of the cells carrying the *IME2-st* allele displayed mitochondrial detachment (Figure 3E and 3F). The remainder had a typical, prophase I mitochondrial morphology. The basis of this heterogeneity is unclear at the moment, though a similar phenotype of incomplete penetrance has been observed in a previous study using the same allele (Berchowitz et al., 2013), potentially suggesting cell-to-cell variation in kinase activity. Nonetheless, these results demonstrate that Ime2 is sufficient to promote mitochondrial detachment and that the requirement of Ime2 for Ndt80 activation and meiotic divisions can be uncoupled from its role in mediating mitochondrial reorganization.

### The mitochondria-plasma membrane tether MECA is phosphorylated in an Ime2-dependent manner

How does Ime2 trigger mitochondrial detachment? A simple model would be that Ime2 inhibits the activity of a factor that normally connects mitochondria to the cell cortex. Since Ime2 is a protein kinase, such a factor could be an Ime2 substrate. A clear candidate was the mitochondria-ER-cortex anchor, MECA (Cerveny et al., 2007; Klecker et al., 2013; Lackner et al., 2013). In mitotic cells, MECA tethers mitochondria to the plasma membrane by forming large assemblies at the contact sites between the two membranes (Cerveny et al., 2007; Klecker et al., 2013; Lackner et al., 2013). MECA has two known subunits, Num1 and Mdm36 (Lackner et al., 2013; Ping et al., 2016). While Num1 can directly bind to lipids on both the outer mitochondrial membrane and plasma membrane, Mdm36 helps Num1 assemble into clusters at the membrane contact sites (Lackner et al., 2013; Ping et al., 2016). Interestingly, in the absence of MECA, mitochondria are constitutively detached from the plasma membrane in vegetative cells (Cerveny et al., 2007; Klecker et al., 2013; Lackner et al., 2013), highly reminiscent of the mitochondrial detachment that naturally occurs as part of meiotic differentiation.

In order to test if Ime2 phosphorylates MECA, we first isolated recombinant Mdm36 and performed an *in vitro* kinase assay with constitutively active Ime2 (Ime2-st) purified from yeast. We found that Mdm36 was phosphorylated only in the presence of Ime2, suggesting that it is a direct substrate for the kinase (Figure 4A and 4B). To analyze Num1, we immunoprecipitated Num1-3V5 or Num1-GFP from vegetative cells and then performed a similar *in vitro* kinase assay. Similar to Mdm36, we observed Ime2-dependent phosphorylation of Num1 (Figure 4C). Together, these results demonstrate that Ime2 can phosphorylate both Num1 and Mdm36 *in vitro.*

**Figure 4.**
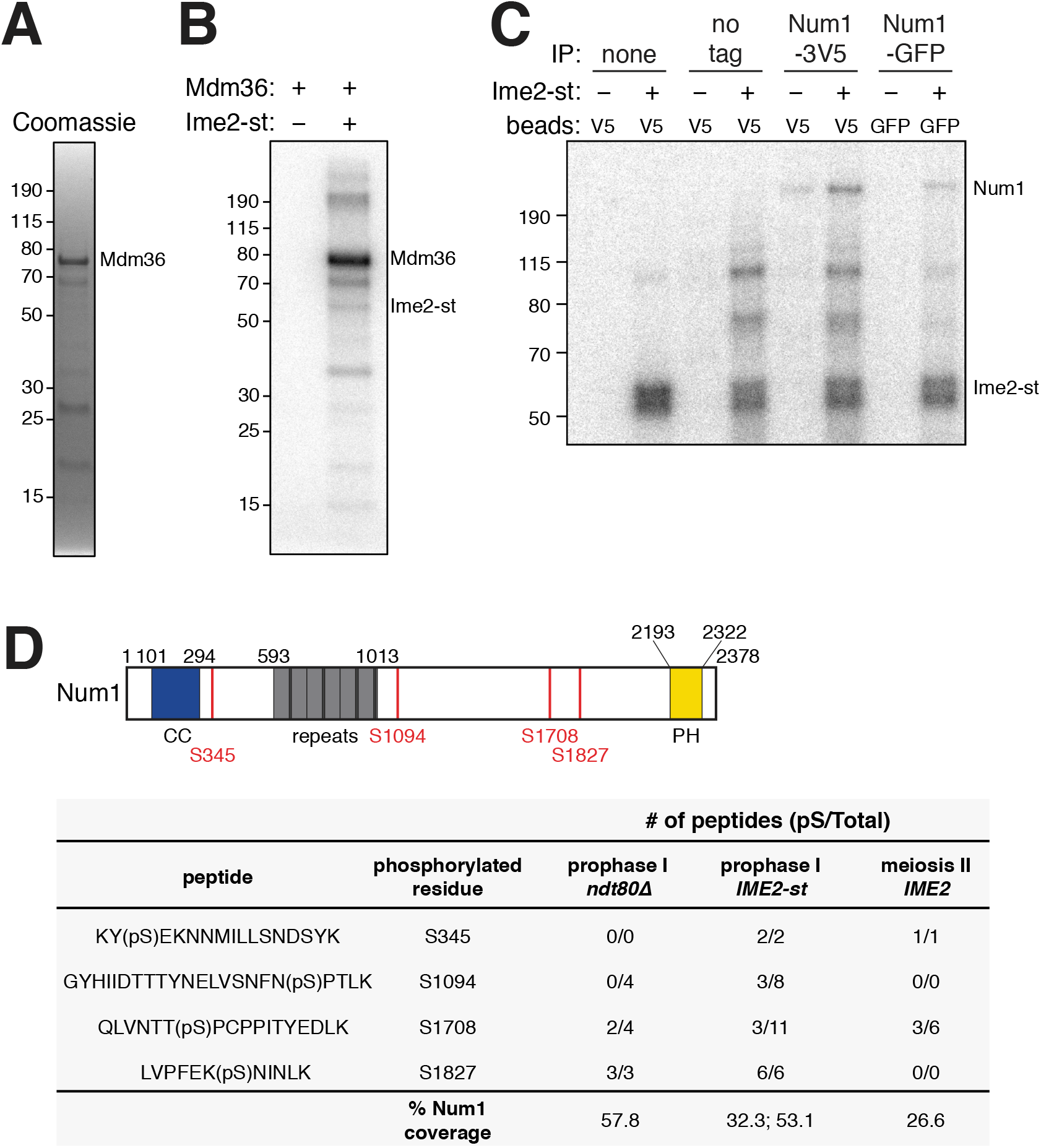
Phosphorylation of Num1 and Mdm36 *in vitro* and *in vivo.* **A**. Coomassie-stained gel of 6xHis-T7-Mdm36 purified from *E. coli,* used in panel B. **B**. *in vitro* kinase assays containing γ-^32^P ATP, 1 *μ*g recombinant Mdm36, and 3 pmol of Ime2-st purified from yeast or no-kinase control. Reactions were incubated for 15 min at room temperature, then analyzed by SDS-PAGE and autoradiography. **C**. *in vitro* kinase assay prepared as in panel B, but using on-bead substrate immunoprecipitated from vegetative cells using agarose beads conjugated with an anti-V5 or anti-GFP antibody. Untagged lysate (“no tag”) was purified from wild type (UB15), Num1-3V5 from strain UB12017, and Num1-GFP from UB15517. Beads never incubated with lysate (“none”) were also examined. **D**. Top: Diagram of Num1 domain structure in the SK1 background and *in vivo* phosphorylation sites detected by mass spectrometry (red lines and text). Black numbers refer to amino acid positions that define the domain boundaries. The Num1 amino acid sequence in SK1 differs from the S288C reference genome (Yue et al., 2017). Num1 (SK1) contains six copies of a 64-amino acid repeat, with some repeats separated by a spacer sequence (LEKEVEQ), and an overall length of 2378 amino acids (271 kDa). CC, coiled coil domain; PH, Pleckstrin homology domain. Bottom: Num1 phosphopeptides detected by LC-MS/MS from Num1-3V5 denaturing immunoprecipitation (see Methods). Phosphoserine is indicated as (pS). The number of phosphopeptides and total peptides (phosphorylated and unmodified) recovered as well as the overall sequence coverage of Num1 is shown. Num1-3V5 was isolated from *ndt80Δ* negative control cells (UB17332; prophase I *ndt80Δ*) and *ndt80Δ IME2-st* cells (UB16660; prophase I *IME2-st*) after 5 h in SPO medium. Num1-3V5 was isolated from *pGAL-NDT80 GAL4.ER* cells (UB12402; meiosis II *IME2*) harvested at 7.5 h, having been released from prophase arrest by the addition of 1 *μ*M β-estradiol at 5 h. Peptide quantifications were pooled from two experiments for the prophase I *IME2-st* sample, with Num1 sequence coverage for each experiment shown.

We further found that Ime2 regulates MECA phosphorylation *in vivo* (Figure 4D). We isolated Num1 from prophase I-arrested cells (*ndt80Δ*) that expressed either wild-type Ime2 or the hyperactive Ime2-st, as well as from wild-type cells enriched in meiosis II, and used label-free mass spectrometry to map phosphorylation sites. This approach led to the identification of four phosphorylated serine residues in Num1, two of which were present only when Ime2 activity was high (Figure 4D). We conclude that Num1, the major subunit of MECA, is phosphorylated in an Ime2-dependent manner during meiosis.

### MECA undergoes dynamic changes in meiosis

To investigate a possible role of MECA in regulating mitochondrial dynamics during meiosis, we first examined the localization of Num1 and Mdm36. Similar to previous studies in vegetative cells (Cerveny et al., 2007; Klecker et al., 2013; Kraft and Lackner, 2017; Lackner et al., 2013; Ping et al., 2016; Tang et al., 2012), Num1 and Mdm36 formed prominent clusters at the cell cortex prior to meiosis II (Figure 5A); these foci represent contact sites between mitochondria and the plasma membrane. In contrast, meiosis II cells were devoid of Num1 and Mdm36 puncta. We further characterized MECA dynamics in meiosis by monitoring Num1-GFP localization in a strain carrying mitoBFP and Htb1-mCherry. In premeiotic and prophase I cells, we observed many bright Num1-GFP puncta, with mitochondria tethered at the plasma membrane (Figure 5B). In contrast, meiosis II cells were largely devoid of the puncta, and mitochondria were detached (Figure 5B). The number of Num1 foci and total cellular fluorescence were significantly reduced in meiosis II cells compared to cells from earlier stages (Figure 5C and 5D). Furthermore, the disappearance of Num1 foci was dependent on Ndt80 expression (Figure 5B-D), but not on APC^Cdc20^ (Figure S2A), consistent with our previous observations of mitochondrial behavior (Figure 2E and Figure 3A). These results strongly indicate that the timely detachment of mitochondria from the plasma membrane is driven by developmentally regulated changes in the MECA complex.

**Figure 5.**
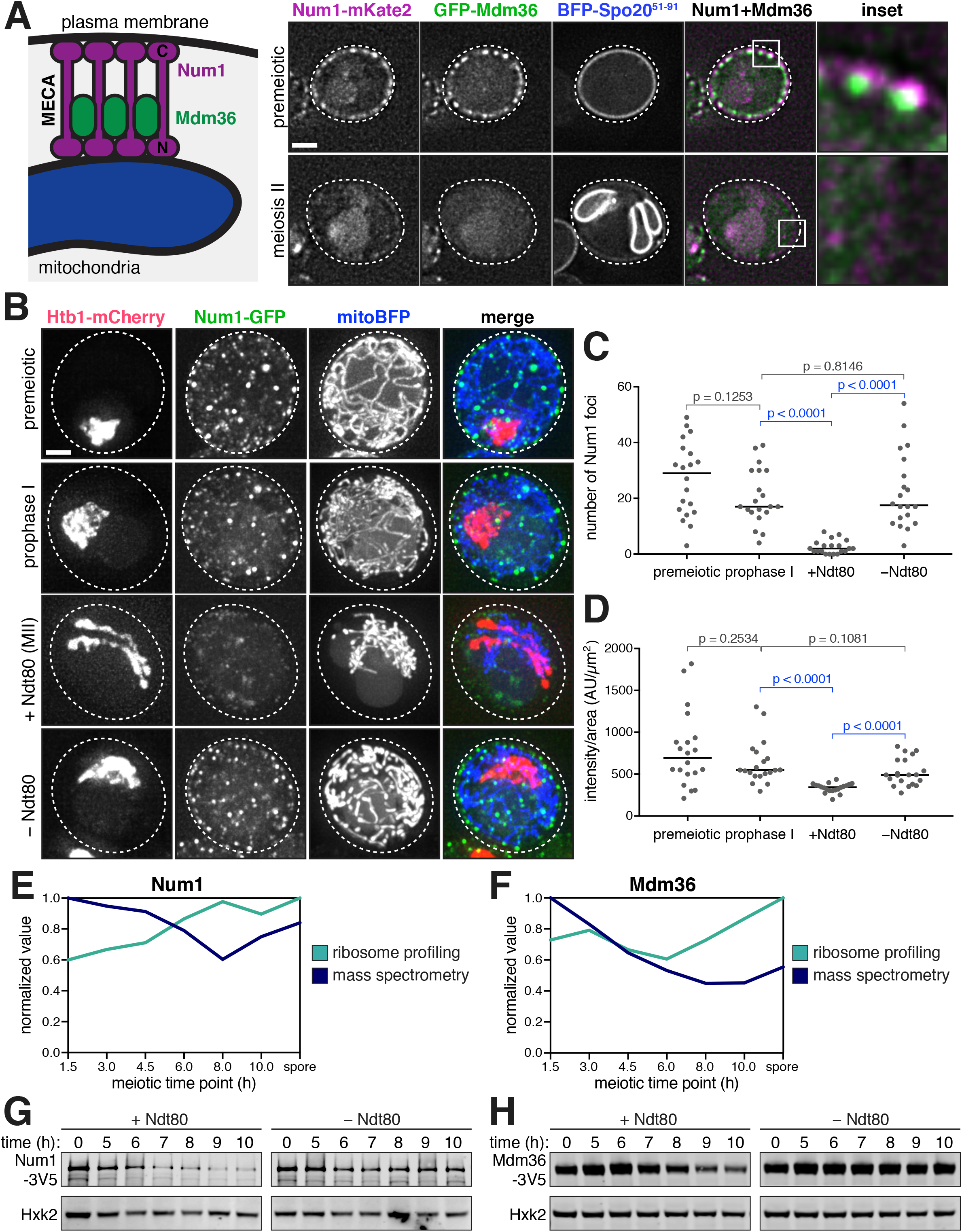
MECA is dynamically regulated in meiosis. **A**. Left: Diagram of the mitochondria-plasma membrane contact site. MECA contains two known subunits, Num1 and Mdm36, and is responsible for tethering mitochondria to the plasma membrane. Right: Localization of MECA subunits, Num1-mKate2 and GFP-Mdm36, in live premeiotic and meiosis II cells (UB16677). Single planes in the z-axis are shown. Meiotic stage was determined by the prospore membrane marker BFP-Spo20^51-91^. **B**. Maximum intensity projections of live *pGAL-NDT80 GAL4.ER* cells (UB15124) showing MECA localization (Num1-GFP), mitochondrial localization (mitoBFP), and nuclei (Htb1-mCherry). Cells were imaged at time of transfer to SPO medium (premeiotic) and 5 h (prophase I). Then, at 5 h, the culture was split and treated with 1 *μ*M β-estradiol (+Ndt80) or ethanol vehicle control (-Ndt80). Cells from the split cultures were imaged at 8 h. **C-D**. Quantifications of the experiment in panel B. Analysis of the +Ndt80 sample was restricted to meiosis II stage cells. *n* = 20 cells per group. Medians are plotted as horizontal lines. The Mann-Whitney nonparametric test was used to test statistical significance. **C**. Number of Num1-GFP foci per cell (see Methods). **D**. Whole-cell Num1-GFP fluorescence quantification from maximum intensity projection (AU, arbitrary units), normalized to cell area. **E-F**. Quantifications of Num1 and Mdm36 steady-state protein levels (mass spectrometry) and synthesis (ribosome profiling) during meiosis from a published dataset (Cheng et al., 2018). **G-H**. Immunoblots of MECA subunits in *pGAL-NDT80 GAL4.ER* strains. Strains were induced to sporulate for 5 h, then flasks were split and treated with 1 *μ*M β-estradiol (+Ndt80) or ethanol vehicle control (-Ndt80). Hxk2 serves as a loading control. **G**. Blot for Num1-3V5 (UB12402). **H**. Blot for Mdm36-3V5 (UB13851). Scale bars, 2 *μ*m.

In a matched ribosome profiling and quantitative mass spectrometry dataset generated from a yeast meiotic time course (Cheng et al., 2018), both Num1 and Mdm36 exhibit dynamic regulation during meiosis. Namely, as assessed by quantitative mass spectrometry, the protein levels of Num1 and Mdm36 decrease as meiosis progresses, reaching their minima at 8 h (Figure 5E and 5F). We confirmed the decline in protein levels by immunoblotting with strains expressing epitope tagged Num1-3V5 or Mdm36-3V5 from their endogenous loci (Figure 5G and 5H, Figure S3). In each case, protein decline occurred in an Ndt80-dependent manner (Figure 5G and 5H), suggesting that Num1 and Mdm36 protein levels are actively regulated.

The decline in Num1 and Mdm36 protein levels is not accompanied by a decrease in ribosome footprint density (Figure 5E and 5F), indicating that the abundance of Num1 and Mdm36 cannot be easily explained by regulation at the level of protein synthesis. Instead, we reasoned that Num1 and Mdm36 are actively degraded. To test whether the reduction in Num1 and Mdm36 protein levels requires the proteasome, a major conduit for protein degradation, we treated prophase I-arrested cells with the proteasome inhibitor MG-132 and simultaneously released them from the Ndt80-block. Upon MG-132 treatment, Num1 protein levels failed to decline to the extent seen in the control (Figure 6A). However, Mdm36 levels continued to decrease (Figure 6B). To assess proteasome-dependence by an independent method, we utilized a hypomorphic allele of a 26S proteasome lid subunit, *rpn6-1* (Isono et al., 2005). Consistent with the MG-132 data, in *rpn6-1* cells, the protein levels of Num1, but not Mdm36, were stabilized throughout meiosis (Figure S2B and S2C), indicating proteasome-dependence for Num1 degradation. To address the possibility that Mdm36 might instead be degraded by autophagy, we utilized the GFP-Mdm36 allele. Since GFP is relatively resistant to vacuolar degradation, autophagic degradation of the tagged protein leads to the accumulation of free GFP in the vacuole (Kanki and Klionsky, 2008). We observed vacuolar GFP signal during meiosis II in cells carrying GFP-Mdm36 by fluorescence microscopy (Figure 6C) and free GFP by immunoblotting (Figure 6D and 6E), consistent with autophagy-dependent turnover of Mdm36. Altogether, these analyses reveal that programmed destruction of Num1 and Mdm36 causes the dissolution of MECA assemblies in meiosis II.

**Figure 6.**
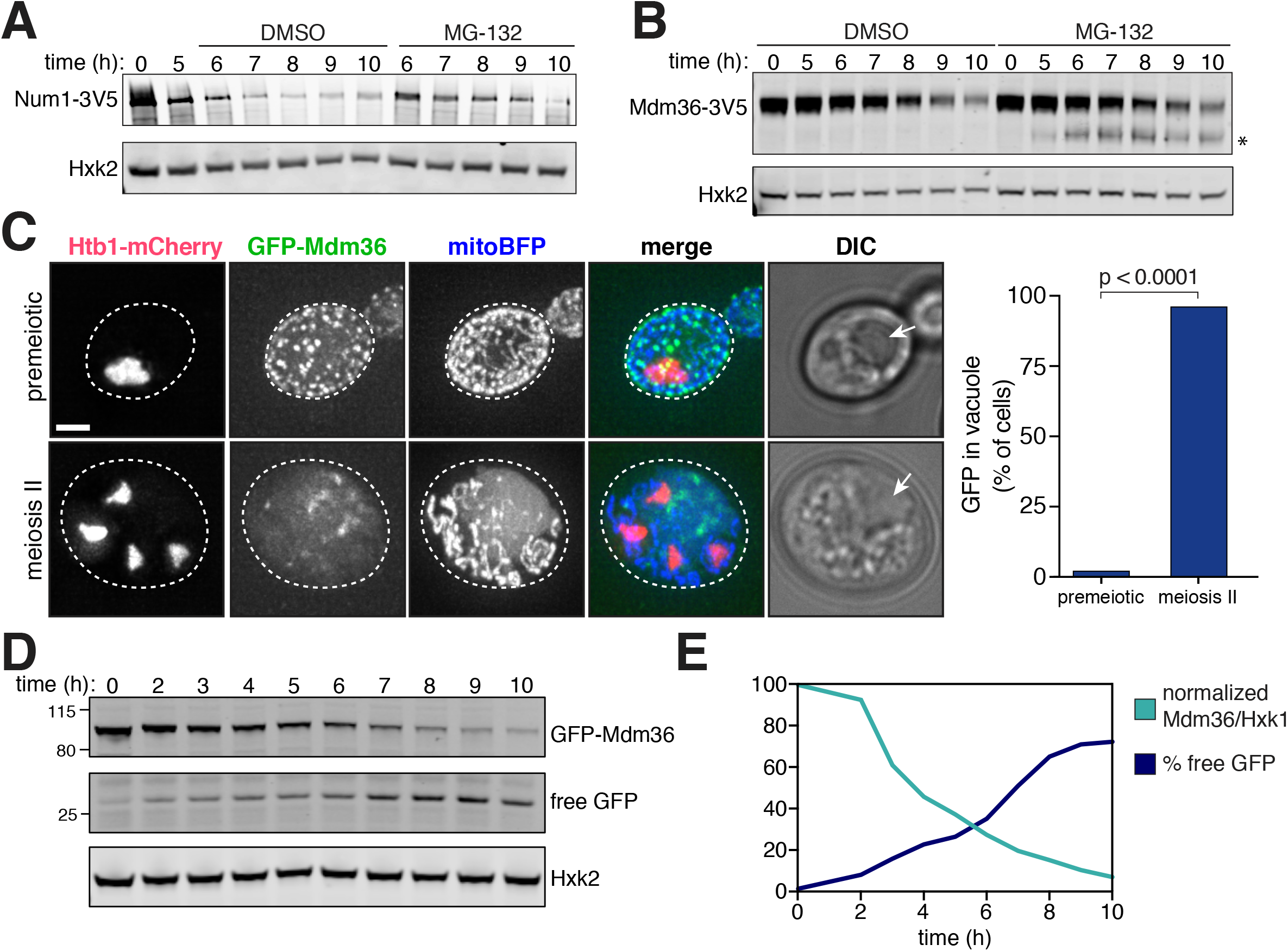
MECA is destroyed by the proteasome and autophagy. **A-B**. Proteasome inhibition during meiosis by treatment with MG-132, compared to a DMSO vehicle control. Flasks were split at 5 h, with one half treated with 100 *μ*M MG-132 and the other half with a DMSO vehicle control. **A**. Immunoblot of Num1-3V5 in a *pGAL-NDT80 GAL4.ER* synchronous meiosis (UB13245), where 1 *μ*M β-estradiol was added to the flasks at 5 h. **B**. Immunoblot of Mdm36-3V5 (UB16324). The asterisk indicates a band of unknown identity. **C**. Left: Maximum intensity projections and DIC images of live cells (UB16683) showing localization of GFP-Mdm36 in premeiotic and meiosis II cells. Arrows indicate the position of the vacuole. Right: Quantification of the frequency of vacuolar GFP signal observed in premeiotic and meiosis II cells. Fisher’s exact test p < 0.0001 (n = 50 per stage). **D**. Immunoblot autophagy assay of GFP-Mdm36 (UB16326). **E**. Quantification of the blot in panel D. The total level of full-length protein (normalized Mdm36/Hxk2) was calculated as the intensity of the GFP-Mdm36 band divided by the Hxk2 loading control band. Percentages of the maximum value are plotted. The % free GFP value was calculated as the intensity of the free GFP band divided by the summed intensities of the free GFP and GFP-Mdm36 bands. Scale bars, 2 *μ*m.

### Ime2 induces mitochondrial detachment by promoting MECA degradation

What is the relationship between Ime2 and MECA in mediating mitochondrial detachment? We posited that Ime2 interferes with MECA function during meiosis II by promoting its destruction, thereby triggering mitochondrial detachment. In order to test this hypothesis, we first examined whether Ime2 regulates MECA stability by measuring Num1 and Mdm36 abundance in the *ime2-as1* mutant background. Treatment with 1-NA-PP1 at the time of Ndt80 block-release attenuated the degradation of Num1 and Mdm36 that was observed in wild-type cells (Figure 7A and 7B). Monitoring MECA assemblies in single cells yielded similar results (Figure 7C). Num1-GFP clusters persisted at later meiotic time points upon Ime2 inhibition (Figure 7D and 7E). In these cells, mitochondria remained attached to the plasma membrane, presumably through the persistent contact sites. The simplest interpretation of these data is that Ime2-dependent regulation leads to Num1 and Mdm36 degradation, which in turn causes MECA disassembly and mitochondrial detachment in anaphase II.

**Figure 7.**
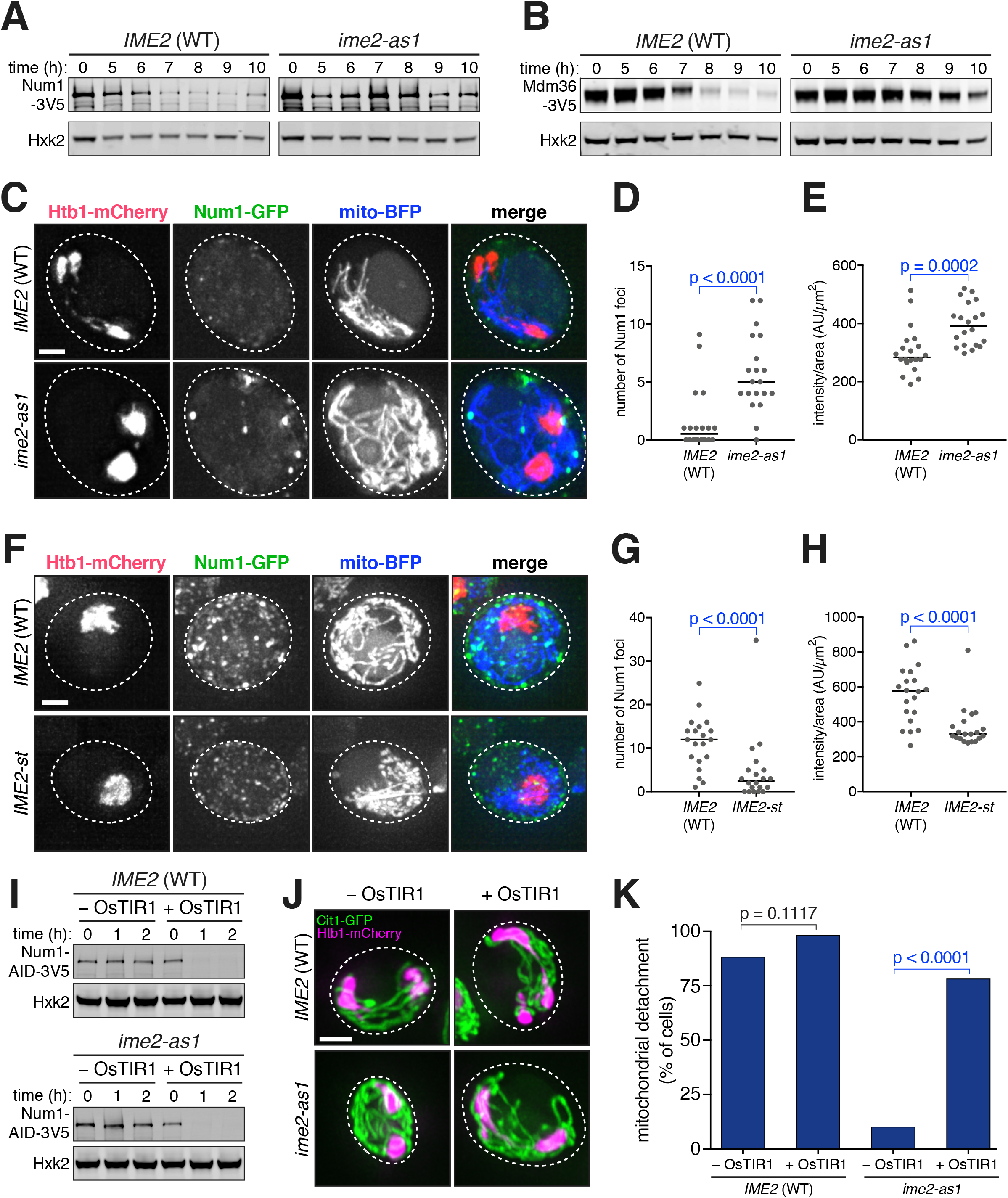
Ime2 regulates MECA function in meiosis. **A-B**. Immunoblots of MECA subunits in *pGAL-NDT80 GAL4.ER.* Strains were induced to sporulate, then 5 h later released from prophase I arrest by the addition of 1 *μ*M β-estradiol to induce *NDT80.* Simultaneously, 20 *μ*M 1-NA-PP1 was added to inhibit *ime2-as1,* with drug-insensitive *IME2* as a wild-type control. Hxk2, loading control. A. Num1-3V5 immunoblot in wild type (UB12402) and *ime2-as1* (UB12403). **B**. Mdm36-3V5 immunoblot in wild type (UB13851) and *ime2-as1* (UB14546). **C**. Maximum intensity projections of live *pGAL-NDT80 GAL4.ER* cells containing wild-type *IME2* (UB15124) or *ime2-as1* (UB16047). Cells were induced to sporulate, then treated with 1 *μ*M β-estradiol and 20 *μ*M 1-NA-PP1 at 5 h. Images were acquired at 8 h. Dashed lines indicate cell boundaries. Mitochondria, mitoBFP. Nuclei, Htb1-mCherry. **D-E**. Quantifications for the experiment in panel C. Analysis was restricted to cells that had entered the meiotic divisions. *n* = 20 cells per group. Medians are plotted as horizontal lines. The Mann-Whitney nonparametric test was used to test statistical significance. **D**. Number of Num1-GFP foci per cell (see Methods). **E**. Whole-cell Num1-GFP fluorescence quantification from maximum intensity projection, normalized to cell area. **F**. Maximum intensity projections of live wild-type *IME2 ndt80Δ* cells (UB16806) or *IME2-st ndt80Δ* cells (UB16808) arrested in prophase I for 5 h. Cells express the same cellular markers as in panel C. **G-H**. Quantifications of the experiment in panel **F**. *n* = 20 cells per group. Medians are plotted as horizontal lines. The Mann-Whitney nonparametric test was used to test statistical significance. **G**. Number of Num1-GFP foci per cell (see Methods). **H**. Wholecell Num1-GFP fluorescence quantification from maximum intensity projection, normalized to cell area. **I-K**. *pGAL-NDT80 GAL4.ER* cells without OsTIR1 (UB17552) and with *pCUP1-OsTIR1* (UB17548) as well as *pGAL-NDT80 GAL4.ER ime2-as1* cells without OsTIR1 (UB17554) and with *pCUP1-OsTIR1* (UB17550) were induced to sporulate. Then, 1 *μ*M β-estradiol and 20 *μ*M 1-NA-PP1 were added to the cultures at 5 h. At 6.5 h, 50 *μ*M CuSO_4_ and 500 *μ*M 3-indoleacetic acid (auxin) were added. **I**. Immunoblot showing Num1-AID-3V5 depletion. Hxk2, loading control. Time indicates hours after addition of auxin and CuSO_4_ at 6.5 h, a time at which Num1 protein level is already reduced. By band quantification normalized to loading, Num1-AID-3V5 is 2.3x higher in level in *ime2-as1* (no OsTIR1) compared to the wild-type control (no OsTIR1). **J**. Maximum intensity projections of cells fixed 2 h after auxin and CuSO_4_ addition. Mitochondria, Cit1-GFP. Nuclei, Htb1-mCherry. **K**. Quantification of the frequency of mitochondrial detachment among cells fixed 2 h after auxin and CuSO**4** addition. Analysis was restricted to cells that had entered the meiotic divisions. Fisher’s exact test p = 0.1117 (WT –OsTIR1 vs. WT +OsTIR1) and p < 0.0001 (*ime2-as1* –OsTIR1 vs. *ime2-as1* +OsTIR1). *n* = 50 cells per genotype. Scale bar, 2 *μ*m.

To further test the involvement of Ime2 in MECA destruction, we examined the impact of expressing the hyperactive Ime2-st kinase on MECA levels in prophase I-arrested cells. Both the number of Num1 foci and total cellular fluorescence were significantly reduced upon *IME2-st* expression (Figure 7F-H). Our results indicate that elevated Ime2 activity in prophase I is sufficient to trigger premature Num1 degradation.

If Ime2 acts through MECA to promote mitochondrial detachment, then removal of MECA should restore normal mitochondrial dynamics to Ime2-inhibited cells. In order to test this hypothesis, we engineered a version of Num1 that can be degraded in an Ime2-independent manner (Figure 7I), using the auxin-inducible degron system (Nishimura et al., 2009). We found that forced destruction of Num1 rescued the mitochondrial detachment defect observed in *ime2-as1* cells (Figure 7J and 7K), highlighting Num1 as the key Ime2 target that is responsible for altering mitochondrial distribution during meiosis.

## DISCUSSION

In this study, we have shown that organelle morphogenesis during cellular differentiation can be accomplished by programmed removal of organelle tethers. The mitochondria-ER-cortex anchor (MECA), which normally localizes to and maintains contact sites between mitochondria and the plasma membrane, is inactivated in meiosis II. As a consequence, mitochondria detach from the plasma membrane in a temporally coordinated manner. Mitochondrial detachment is regulated by the meiosis-specific kinase Ime2, which phosphorylates MECA and promotes its destruction (Figure 8). Altogether, our study defines a key mechanism that coordinates mitochondrial dynamics with meiotic progression and demonstrates that organelle remodeling can be mediated by posttranslational regulation of organelle tethers.

**Figure 8.**
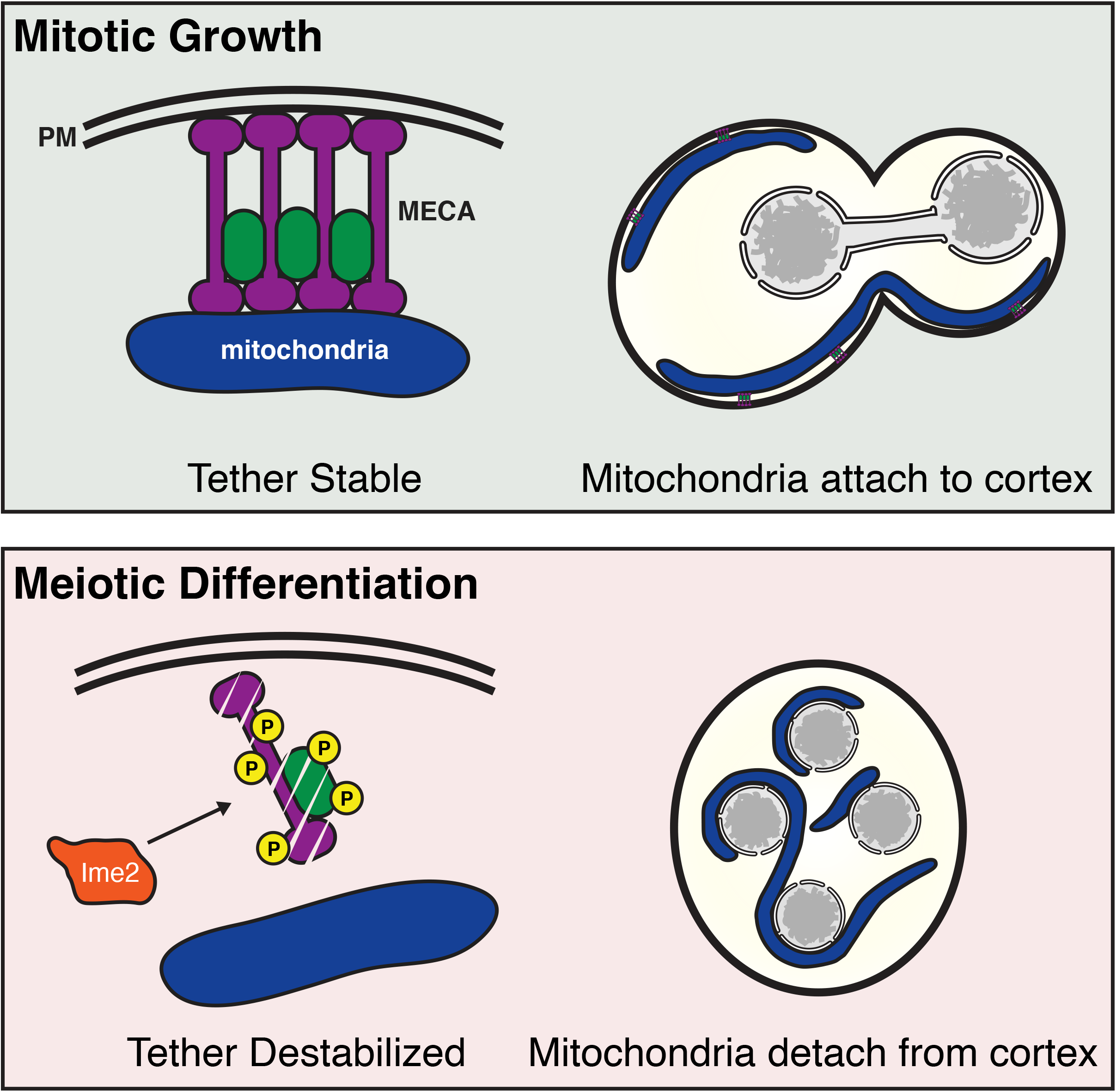
Mitochondrial inheritance in mitosis and meiosis. In mitosis, mitochondria remain associated with the cell cortex due to the mitochondria-plasma membrane anchoring activity of MECA. In meiosis, mitochondrial organization is remodeled: mitochondria detach from the plasma membrane and are transmitted to spores. This meiosis-specific mitochondrial remodeling is due to inhibition of MECA by Ime2. As a result of Ime2-dependent phosphorylation, MECA is destroyed and mitochondrial tethering is lost. PM, plasma membrane.

### Regulated destruction of an organelle tether acutely changes intracellular organization

In vegetative cells, the timing of MECA assembly in the daughter cell is determined by the timing of mitochondrial transport. If transport of mitochondria into the bud is disrupted, MECA assemblies fail to form (Kraft and Lackner, 2017). It is not clear, however, whether vegetative cells regulate the disassembly of MECA in any meaningful way. Once MECA is assembled, it is a very stable anchor: by FRAP, no appreciable recovery was observed for over 20 min (Kraft and Lackner, 2017). Thus, it appears that MECA is a source of stability for an organelle that is otherwise highly dynamic in its architecture (Friedman and Nunnari, 2014; Mishra and Chan, 2014; Westermann, 2014).

During meiosis, MECA undergoes temporally coordinated disassembly. The timing of MECA destruction is determined by the level of Ime2 activity, which in turn is controlled by the meiotic transcription factor Ndt80. Our study demonstrates that an organelle tether can be developmentally regulated and reveals how this regulation could impact organelle remodeling. A similar principle might apply to other organelles and different developmental contexts.

In meiotic cells, the cortical ER detaches from the plasma membrane in a manner highly reminiscent of mitochondrial detachment (Suda et al., 2007). MECA destruction probably does not explain this behavior. Although ER has been observed in association with MECA by light microscopy and co-immunoprecipitation (Lackner et al., 2013), electron microscopy analysis did not reveal a significant association (Klecker et al., 2013). Additionally, in neither case was it suggested that MECA acts to tether the ER to the plasma membrane. A different study found that six proteins, none of which are MECA subunits, redundantly establish ER-plasma membrane contacts (Manford et al., 2012). Based on the finding that removal of any one tether or class of tethers was insufficient for detachment of cortical ER from the plasma membrane, it would appear that meiotic cells have to simultaneously target multiple tethers in order to disrupt ER-plasma membrane contacts. Alternatively, changes intrinsic to the ER or plasma membrane, such as protein or lipid composition, might explain these phenomena. Interestingly, the plasma membrane pool of the lipid phosphatidylinositol 4,5-bisphosphate (PIP2) is depleted in late meiosis, instead accumulating on prospore membranes (Rudge et al., 2004).

### Ime2 is a key regulator of mitochondrial dynamics in meiosis

Our studies reveal a distinct and unanticipated function for Ime2 during meiosis II: regulation of the mitochondria-plasma membrane tether, MECA. Our data support a model in which Ime2 triggers MECA destruction by promoting the phosphorylation of its subunits, thereby causing acute changes in mitochondrial organization (Figure 8). Several observations are consistent with this model: First, Ime2 phosphorylates both MECA subunits *in vitro.* Second, inactivation of Ime2 causes stabilization of MECA subunits, persistence of MECA clusters, and retention of contacts between mitochondria and the plasma membrane throughout meiosis. Third, expression of a hyperactive *IME2* allele in prophase I leads to Num1 phosphorylation and results in the premature disassembly of MECA and untimely mitochondrial detachment. Finally, induced degradation of Num1 rescues the mitochondrial detachment defect that occurs in *IME2* inactivated cells. Our data, however, do not rule out the possibility that Ime2 can also influence MECA in an indirect manner, for instance through its effect on other, yet to be identified, MECA regulators.

Ime2-dependent destruction of MECA shares similarities with another critical meiotic event: clearance of the translational repressor Rim4. Rim4 assembles into amyloid-like aggregates, which are thought to sequester a group of mRNAs away from ribosomes by binding to their 5’ untranslated regions (Berchowitz et al., 2013; Berchowitz et al., 2015; Carpenter et al., 2018). Degradation of Rim4 during meiosis II relieves the translational repression of its targets. Similar to MECA, high Ime2 kinase activity is both necessary and sufficient to disassemble Rim4 aggregates and promote their degradation (Berchowitz et al., 2013; Berchowitz et al., 2015; Carpenter et al., 2018). Rim4 contains a total of 114 serine and threonine residues (S/Ts). Although the initial mass spectrometry suggested the existence of a single Ime2-dependent phosphorylation site, subsequent work identified 39 additional phosphorylation sites. Clearance of Rim4 assemblies is governed by multi-site phosphorylation, with at least 36 S/Ts required for its degradation. Importantly, a threshold amount of phosphorylation, rather than modification of critical residues, is necessary for Rim4 clearance (Carpenter et al., 2018). By comparison, Num1 contains 356 S/Ts. Thus far, mass spectroscopy identified four phosphorylated residues in Num1, two of which appear to be Ime2-dependent. However, this number is likely to be an underestimate, because the peptide coverage for Num1 was less than 60% in our immunoprecipitation-mass spectrometry (IP-MS) analysis. Moreover, unlike the Rim4 IP-MS, our experiments did not include a phospho-peptide enrichment step due to the low expression level of Num1. Adding further to the complexity is the observation that the second MECA subunit, Mdm36, also appears to be phosphorylated by Ime2. Therefore, MECA control by Ime2 is likely to be highly complex. More thorough analysis will be required to elucidate how phosphorylation affects MECA stability, mitochondrial organization, and inheritance.

### Mitochondrial inheritance during gametogenesis

How does mitochondrial detachment lead to meiosis-specific inheritance of the organelle? In budding yeast, mitochondria exhibit four distinct behaviors during meiotic differentiation: (i) abrupt detachment from the mother cell plasma membrane, followed by (ii) extensive contacts with the gamete nuclei, (iii) limited inheritance, and (iv) programmed elimination. Previous electron microscopy data suggested that only about half of the starting mitochondrial population is inherited by the four gametes (Brewer and Fangman, 1980). The remaining mitochondria are eliminated by mega-autophagy that commences at the end of gametogenesis (Eastwood et al., 2012; Eastwood and Meneghini, 2015). It will be interesting to determine if the two populations of mitochondria, namely the inherited and discarded, are different from one another and whether quality control pathways exist to selectively transmit healthier mitochondria to gametes. Perhaps the ability of mitochondria to form direct contact sites with the nuclear envelope is part of this selection (note the extent of mitochondria-nucleus interaction in Figure 2). Regardless, we propose that the contact sites between mitochondria and nuclear envelope ensure that the mitochondrial genome is inherited during yeast gametogenesis. We further posit that the regulated detachment of mitochondria from the progenitor cell plasma membrane in meiosis II is the first step towards mitochondrial segregation into gametes. Identifying the molecular nature of the mito-nuclear contact sites and their regulation will enhance our understanding of mitochondrial inheritance during meiotic differentiation.

Gametogenesis-specific changes to mitochondrial architecture and inheritance are ubiquitous in metazoan germ cells. For example, primary oocytes of animals contain a unique structure known as the Balbiani body that assembles adjacent to the nucleus (Kloc et al., 2004). The Balbiani body houses a collection of organelles, including mitochondria as well as protein–RNA inclusions, and facilitates their segregation. In *Drosophila* oogenesis, mitochondria are transported between cells, from nurse cells to the oocyte, via a polarized microtubule network that passes through ring canals (Cox and Spradling, 2003). Later, mitochondria are actively tethered to the actin cytoskeleton at the posterior of the oocyte, in proximity to the pole cells that give rise to the germline. Importantly, in the absence of tethering, mtDNA transmission is compromised (Hurd et al., 2016). Although sperm contain mitochondria to meet metabolic demands, they do not transmit genetic information to the zygote. In *Drosophila,* mtDNA is actively destroyed during spermatogenesis (DeLuca and O’Farrell, 2012). In mice, sperm mitochondria are delivered to the zygote, but are depolarized, unable to fuse to maternal mitochondria, and are specifically eliminated by mitophagy (Rojansky et al., 2016). Clearly, mitochondria undergo a plethora of changes during gametogenesis. Understanding the molecular basis of meiotic specializations to mitochondria is important not only to enhance our understanding of the organelle’s physiology but also for its potential impact on human disease. It is widely observed that mitochondrial functions decline with age, yet gametogenesis, at least in budding yeast and *C. elegans,* eliminates age-induced cellular damage (Bohnert and Kenyon, 2017; Goudeau and Aguilaniu, 2010; Unal et al., 2011). Therefore, studying mitochondria in the context of gametogenesis could aid in the development of strategies to counteract mitochondrial dysfunction and disease.

## METHODS

### Yeast strains and plasmids

All yeast strains used in this study are derivatives of SK1, except for B114, and are described in Table S1. The Ndt80 block-release system and associated strains were described previously (Benjamin et al., 2003; Carlile and Amon, 2008). The PP1 analog sensitive kinase alleles *cdc28-as1* (F88G) (Bishop et al., 2000) and *ime2-as1* (M146G) (Benjamin et al., 2003) were previously described. The *IME2-st* allele was described previously (Sia and Mitchell, 1995). Meiotic-null alleles generated by promoter replacement by *pCLB2* were described previously for *CDC5* and *CDC20* (Lee and Amon, 2003). The *rpn6-1* (*435Y) allele was described previously (Isono et al., 2005), and the His3MX6-marked SK1 version of the strain was described previously (Carpenter et al., 2018). The auxin-inducible degron system was described previously (Nishimura et al., 2009). In this study, we used TIR1 from *Oryza sativa* under regulation of a copper-inducible promoter, *pCUP1.* The *pCUP1-OsTIR1* construct was cloned into the *HIS3* single integration vector pNH603 (Youk and Lim, 2014), but modified to remove homology to the *DED1* locus (a gift from Leon Chan). The Num1-AID allele carried a C-terminal IAA7 degron followed by a 3V5 tag. The IAA7-3V5 tagging plasmid was a gift from Leon Chan.

Yeast transformation was performed using the lithium acetate method. C-terminal tagging was performed using a PCR-mediated technique previously described (Janke et al., 2004; Longtine et al., 1998). As some C-terminally tagged alleles of *MDM36* are not functional, we verified functionality of *MDM36-3V5* using an established assay (Figure S3; (Lackner et al., 2013)). We constructed a *LEU2-selectable* GFP(S65T) tagging plasmid by replacing the selectable marker in pFA6a-GFP(S65T)-His3MX6 with *Candida glabrata LEU2 (cgLEU2),* amplified from pLC605 (a gift from Leon Chan). The 3V5 tagging plasmid was a gift from Vincent Guacci.

To visualize mitochondria, we employed several different strategies. First, we C-terminally tagged *CIT1* with a fluorescent protein, as described (Higuchi-Sanabria et al., 2016), using GFP(S65T) or mCardinal (Chu et al., 2014). The mCardinal yeast tagging plasmid was a gift from Ryo Higuchi-Sanabria. We also expressed the Su9 mitochondrial targeting sequence (MTS) from *Neurospora crassa* fused to yEGFP under regulation of a *pGPD1* promoter, expressed from a pNH603 single integration plasmid (Youk and Lim, 2014), but modified to remove homology to the *DED1* locus (a gift from Leon Chan). Last, we expressed a mitoBFP construct from a pRS424 plasmid modified to carry a KanMX or NatMX marker instead of *TRP1* (termed pRS(2μ)-KanMX or - NatMX). The mitoBFP construct was described as pYES-TagBFP (Murley et al., 2013), and pRS424 was described previously (Christianson et al., 1992).

To visualize the prospore membrane, we fused amino acids 51-91 from Spo20 (Nakanishi et al., 2004) to the C terminus of link-yEGFP (Sheff et al., 2004) or mTagBFP2, under regulation of the *ATG8* promoter (amplified from pRS306-2xyEGFP-ATG8 (Graef et al., 2013)), subcloned into the *LEU2* integrating plasmid pLC605 (a gift from Leon Chan) or a pRS(2μ), drug-selectable plasmid (described above).

To generate the *GFP-MDM36* strain, we used a Cas9-mediated genome editing strategy similar to a described method (Anand et al., 2017). Annealed oligonucleotides encoding the gRNA (5’-GAACACTTACTACTATAGCA-3’) were inserted into a centromeric plasmid carrying a *URA3* marker and *pPGK1-Cas9* (a gift from Gavin Schlissel), generating pUB1395. Then, pUB1395 and a repair template were cotransformed into yeast, the plasmid was lost by streaking cells without selection, and the presence of the yEGFP tag was confirmed by PCR and sequencing.

To purify Mdm36 from *E. coli,* we used an IPTG-inducible expression plasmid described previously as pET22b mod T7prom::H6-T7-Mdm36 (Ping et al., 2016) and provided by Laura Lackner.

### Sporulation

Unless indicated otherwise, cells were induced to sporulate by a traditional starvation synchronization method. At all steps, flasks were shaken at 275 rpm. First, cells were grown in YPD (1% yeast extract, 2% peptone, 2% glucose, 22.4 mg/L uracil, and 80 mg/L tryptophan) for ~24 h at room temperature to reach saturation (OD_600_ ≥ 10). The YPD culture was used to inoculate BYTA medium (1% yeast extract, 2% bacto tryptone, 1% potassium acetate, 50 mM potassium phthalate) to OD_600_ = 0.25 and grown for ~16 h at 30°C to reach OD_600_ ≥ 5. Then, the cells were pelleted, washed with sterile water, and resuspended to a density of OD_600_ = 1.85 in SPO media (0.5% potassium acetate, 0.02% raffinose, pH 7). Cultures were shaken at 30°C for the duration of the experiment. In cases where selection for plasmids was necessary, G418 (Geneticin) or nourseothricin (clonNAT) were added to YPD and BYTA cultures at concentrations of 200 μg/mL and 100 μg/mL, respectively.

In experiments utilizing synchronization by the Ndt80 block-release system, cells carrying the *pGAL-NDT80* and *GAL4.ER* transgenes were induced to sporulate as described above. After 5 h in SPO medium, β-estradiol was added to a final concentration of 1 μM from a 5 mM stock (in ethanol) to induce *NDT80* expression.

### Microscopy

All images were acquired using a DeltaVision Elite wide-field fluorescence microscope (GE Healthcare, Sunnyvale, CA). Time-lapse imaging experiments were performed in an environmental chamber heated to 30°C, with images acquired using a 60X/NA1.42 oil-immersion objective. Cells were maintained on concanavalin A-coated, glass-bottom 96-well plates (Corning). Every 10 min, a stack of 8 Z positions (1 μm step size) was acquired, with mCherry (32% intensity, 25 ms exposure) and FITC (10% intensity, 25 ms exposure) filter sets. All other imaging experiments were acquired using a 100X/NA1.4 oil-immersion objective. Where indicated, cells were fixed at room temperature for 15 min by adding formaldehyde (final concentration of 3.7%) directly to the culture medium. To halt fixation, cells were washed once with 100 mM potassium phosphate, pH 6.4, and stored in 100 mM potassium phosphate pH 7.5 with 1.2 M sorbitol at 4°C. All images were deconvolved in softWoRx software (GE Healthcare) using a 3D iterative constrained deconvolution algorithm (enhanced ratio) with 15 iterations. Linear adjustments to brightness and contrast were made in FIJI.

### Image quantification

Num1-GFP spot number and Num1-GFP fluorescence intensity were measured in FIJI. First, the mean background intensity was measured in a 155 x 155 pixel square containing no cells. This value was then subtracted from each pixel in the image. Next, cells were manually traced, and the Find Maxima function was run to identify spots (noise tolerance = 1500) within the traced region. In addition, the total fluorescence intensity (IntDen) and area were measured for each cell.

### Immunoblotting

To harvest protein, 3.7 OD_600_ equivalents of cells were pelleted and resuspended in 1 mL of 5% trichloroacetic acid (TCA) and incubated at 4°C for ≥10 min. Then, cells were washed with 1 mL of TE50 pH 7.5 (50 mM Tris, 1 mM EDTA), and finally with 1 mL of acetone, then allowed to dry completely. To extract protein, ~100 μL of glass beads and 100 μL of lysis buffer (TE50 pH 7.5, 2.75 mM DTT, 1X cOmplete EDTA-free protease inhibitor cocktail [Roche]) were added to the pellet and shaken using a Mini-Beadbeater-96 (BioSpec). Next, 50 μL of 3X SDS sample buffer (187.5 mM Tris pH 6.8, 6% β-mercaptoethanol, 30% glycerol, 9% SDS, 0.05% bromophenol blue) was added, and the mixture heated for 5 min. We found that recovery of Num1 was enhanced by heating at 50°C rather than boiling.

Proteins were separated by SDS-PAGE using Bolt 4-12% Bis-Tris Plus Gels (Thermo Fisher) and transferred onto membranes. For Num1, we transferred to a 0.45 μm Immobilon-FL PVDF membrane (LI-COR Biosciences, Lincoln, NE) in a Mini-PROTEAN Tetra tank (BioRad) filled with 25 mM Tris, 192 mM glycine, and 9% methanol, run at 180 mA (max 80 V) for 3 h. For all other blots, we transferred to a 0.45 μm nitrocellulose membrane (BioRad) using a semi-dry transfer system (Trans-Blot Turbo) and its supplied transfer buffer (BioRad). Membranes were blocked for 30 min with Odyssey Blocking Buffer (PBS) (LI-COR Biosciences) at room temperature, then incubated overnight at 4°C with primary antibody mixtures diluted in Odyssey Blocking Buffer (PBS) with 0.1% Tween-20. For detection of V5 epitope, we used a mouse anti-V5 antibody (RRID:AB_2556564, R960-25, Thermo Fisher) at a 1:2000 dilution. For detection of GFP, we used a mouse-anti-GFP antibody (RRID:AB_2313808, 632381, Clontech) at a 1:2000 dilution. As a loading control, we used a rabbit anti-hexokinase (Hxk2) antibody (RRID:AB_219918, 100-4159, Rockland, Limerick, PA) at 1:15,000 dilution. For secondary detection, we used an anti-mouse secondary antibody conjugated to IRDye 800CW at a 1:15,000 dilution (RRID:AB_621847, 926–32212, LI-COR Biosciences) and an anti-rabbit antibody conjugated to IRDye 680RD at a 1:15,000 dilution (RRID:AB_10956166, 926–68071, LI-COR Biosciences) in Odyssey Blocking Buffer (PBS) with 0.01% Tween-20. Blots were imaged using an Odyssey CLx scanner (LI-COR Biosciences), and band intensities were quantified using the Image Studio software associated with the scanner.

### Protein purification

Purification of His-tagged Mdm36 was performed as described (Ping et al., 2016) with minor modifications. In brief, 500-mL cultures of Rosetta 2(DE3) *E. coli* (Novagen) bearing expression plasmids and growing in log phase were induced with 250 μM IPTG for 16 h at 18°C. Then, cells were harvested by centrifugation, the pellet resuspended in resuspension buffer (RB, 20 mM Tris pH 8, 500 mM NaCl, 1.89 mM 2-mercaptoethanol), and lysed by 3 freeze-thaw cycles and sonication. Clarified lysates were mixed with 1/7 volume of Ni-NTA agarose (Qiagen) and rotated end-over-end for 1 h at 4°C. Beads were washed in a conical tube with RB, then loaded into a chromatography column and washed with 500 mL of RB + 30 mM imidazole. Protein was eluted with RB + 300 mM imidazole, then dialyzed against 20 mM Tris pH 8, 500 mM NaCl. Last, glycerol was added to 10%, proteins were aliquotted, and flash-frozen in liquid nitrogen. Protein concentration was estimated by A280.

Ime2st kinase was purified from yeast (B114), described previously as yDP159 (Phizicky et al., 2018), with minor modifications. First, a 100-mL culture of SC -Leu with strain B114 was grown overnight with shaking at 30°C. The following day, this culture was used to inoculate 4 L of YEP + 2% glycerol (8 x 500-mL cultures in 2-L baffled flasks) and grown overnight with shaking at 30°C. The following day when the culture reached OD_600_ 1.2, expression of pGAL-IME2st-3xFLAG was induced by the addition of galactose to a final concentration of 2%. (The protein contains amino acids 1-404 of Ime2 fused to a 3x FLAG epitope at the C terminus). After an additional 6 h of growth, cells were harvested by centrifugation at 4°C.

The yeast pellet was then resuspended in 35 mL of lysis buffer (50 mM HEPES pH 7.6, 10% glycerol, 5 mM Mg-Acetate, 1 mM EDTA, 1 mM EGTA, 1 M sorbitol, 0.02% NP-40, 2 mM ATP, 0.5 M KCl, 1X Halt protease and phosphatase inhibitors) and cells were drop-frozen in liquid nitrogen. Frozen cells were lysed under liquid nitrogen in a Waring blender. The resulting powder was thawed and clarified for 1 h at 20,000 rpm at 4°C in a JA-20 rotor. The supernatant was adjusted to 0.3 M KCl and clarified again at 25,000 x g for 15 min at 4°C.

To isolate the tagged protein, lysate was incubated with 1 mL of equilibrated FLAG resin (Sigma) for 2 hours with rotation at 4°C. Then, the lysate/resin was transferred to a gravity flow column (at 4°C) and washed with 20 mL H buffer containing 0.3 M KCl, 0.01% NP-40. Beads were washed again with 10 mL H buffer containing 0.3 M KGlut, 0.01% NP-40. Proteins were eluted with 5 mL H buffer containing 0.15 mg/ml 3xFLAG Peptide and 0.3 M KGlut (5 x 1-mL elutions incubated for 30 min each). The eluate was subsequently concentrated by ultrafiltration (10 kDa MWCO, Vivaspin) and then run through a Superdex 75 column (GE Healthcare) equilibrated with H buffer containing 0.3 M KGlut, 1 mM ATP, 0.01% NP-40 using a ActaPur FPLC (GE Healthcare). Peak Ime2-st fractions were pooled, aliquotted, and stored at −80°C.

### *In vitro* kinase assays

Ime2st kinase and recombinant Mdm36 substrate was purified as described above. Immunoprecipitated substrates were purified as follows. 50 OD_600_ equivalents of cells growing in YPD were pelleted, resuspended in 10 mM Tris pH 7.5, pelleted again, and flash-frozen in liquid nitrogen. Frozen pellets were thawed on ice and resuspended in 300 μL of NP-40 lysis buffer (50 mM Tris pH 7.5, 150 mM NaCl, 1% NP-40, 5% glycerol) containing 1 mM DTT, 1 mM EDTA, 3X cOmplete Ultra Protease Inhibitors without EDTA (Roche), and 1X PhosSTOP phosphatase inhibitors (Roche). Cells were broken on a Mini-Beadbeater-96, and extracts were clarified by centrifugation at 15,000 x g for 15 min. Total protein concentration was determined by Bradford Assay (BioRad). For each IP, 1 mg of total protein was used with either V5 agarose beads (Sigma) or GFP-Trap_A agarose beads (Chromotek). After incubation with extract for 2 h at 4°C, beads were washed 4 times with NP-40 lysis buffer and twice with 25 mM MOPS. Then, beads were split for the kinase assay (+Ime2-st and –Ime2-st).

For the kinase assay, recombinant or on-bead substrate was incubated with 6 μL of HBII (15 mM MOPS, 15 mM MgCl_2_, 5 mM EGTA, 1 mM EDTA, 3X cOmplete Ultra without EDTA, 1X PhosSTOP) for 15 min at room temperature. Then, solution 1 was prepared (1.1 μL of 100 mM ATP, 275 μL of water, 3 μL 6000 Ci/mmol 10 mCi/mL γ-^32^P ATP). The kinase reaction was assembled by adding 5 μL of solution 1 and 2 μL of 1.5 μM Ime2-st to the substrate in HBII, resulting in a final volume of 16 μL in 25 mM MOPS. After 15 min at room temperature, reactions were stopped with SDS sample buffer, heated, and run on a SDS-PAGE gel. The gels were fixed in 10% methanol, 10% acetic acid, dried, and exposed to a phosphor screen. Screens were imaged using a Typhoon scanner (GE Healthcare).

### Denaturing immunoprecipitation and mass spectrometry

To generate denatured protein extracts, cells were first pelleted in multiples of 9.25 OD_600_ equivalents (i.e., 5 mL of SPO culture), then resuspended in 2/5 culture volume of 5% TCA at 4°C and distributed into tubes such that each contained 9.25 OD_600_ equivalents. After incubation overnight at 4°C, cells were pelleted, washed once with acetone, and dried completely. To break pellets, 100 μL of zirconia beads and 150 μL of TE50 pH 7.5, 2.75 mM DTT, 1X PhosSTOP (Roche) and 3X cOmplete Ultra EDTA Free (Roche), were added to each tube. Pellets were disrupted on a Mini-Beadbeater-96. Then, SDS was added to 1%, extracts were denatured by heating at 50°C for 5 min, and NP-40 lysis buffer (50 mM Tris pH 7.5, 150 mM NaCl, 1% NP-40, 5% glycerol) was added, supplemented with 1X PhosSTOP and 3X cOmplete Ultra EDTA Free, to a final volume of 1.5 mL (i.e., diluting SDS to 0.1%). Cleared lysates pooled from 5 tubes were added to V5 agarose beads (Sigma), incubated for 2 h at 4°C with rotation, and then washed twice with each: (1) 50 mM Tris pH 7.5, 1 M NaCl, 1 mM EDTA, 1% NP40; (2) 50 mM Tris pH 7.5, 150 mM NaCl, 10 mM MgCl_2_, 0.05% NP-40, 5% glycerol; and (3) 50 mM Tris pH 7.5, 150 mM NaCl, 10 mM MgCl_2_, 5% glycerol. After washes, 1X SDS sample buffer was added to the beads, and proteins were eluted by heating at 50°C for 5 min. Eluted proteins were separated by SDS-PAGE, then stained using a Colloidal Blue Staining Kit (Invitrogen).

Gel bands containing the desired protein were excised, washed for 20 min in 500 μL of 100 mM NH_4_HCO_3_, then incubated at 50°C for 15 min in 150 μL of 100 mM NH_4_HCO_3_ and 2.8 mM DTT. 10 μL of 100 mM iodoacetamide was then added to the cooled gel band mixtures and incubated for 15 min in the dark at room temperature. Then the gel slice was washed in 500 μL of equal parts 100 mM NH_4_HCO_3_ and acetonitrile with shaking for 20 min. Gel slices were shrunk by soaking in 50 μL of acetonitrile for 15 min. Then the supernatant was removed, and residual solvent was removed in a speed vac. Gel fragments were rehydrated with 10 μL of 25 mM NH_4_HCO_3_ containing sequencing grade modified trypsin (Promega), incubated at room temperature for 15 min, then supplemented with additional trypsin to completely cover the gel slices. Digestion was allowed to continue overnight at 37°C. Then, the supernatant; two washes with 60% acetonitrile, 0.1% formic acid; and one wash with acetonitrile were all combined and dried completely in a speed vac.

Mass spectrometry was performed by the Vincent J. Coates Proteomics/Mass Spectrometry Laboratory at UC Berkeley. mudPIT methods were used in order to achieve good sequence coverage of target proteins in a complex mixture. A nano LC column was packed in a 100 μm inner diameter glass capillary with an emitter tip. The column consisted of 10 cm of Polaris c18 5 μm packing material (Varian), followed by 4 cm of Partisphere 5 SCX (Whatman). The column was loaded by use of a pressure bomb and washed extensively with buffer A (see below). The column was then directly coupled to an electrospray ionization source mounted on a Thermo-Fisher LTQ XL linear ion trap mass spectrometer. An Agilent 1200 HPLC equipped with a split line so as to deliver a flow rate of 300 nL/min was used for chromatography. Peptides were eluted using an 8-step mudPIT procedure (Washburn et al., 2001). Buffer A was 5% acetonitrile, 0.02% heptaflurobutyric acid (HBFA). Buffer B was 80% acetonitrile, 0.02% HBFA. Buffer C was 250 mM ammonium acetate, 5% acetonitrile, 0.02% HBFA. Buffer D was same as buffer C, but with 500 mM ammonium acetate.

Protein identification was performed with Integrated Proteomics Pipeline (IP2, Integrated Proteomics Applications, Inc., San Diego, CA) using ProLuCID/Sequest, DTASelect2, and Census (Cociorva et al., 2007; Park et al., 2008; Tabb et al., 2002; Xu et al., 2015). Tandem mass spectra were extracted into ms1 and ms2 files from raw files using RawExtractor (McDonald et al., 2004). Data were searched against the SK1 sequence of the target protein (Yue et al., 2017) plus the yeast database supplemented with sequences of common contaminants and concatenated to a decoy database in which the sequence for each entry in the original database was reversed (Peng et al., 2003). LTQ data was searched with 3000.0 milli-amu precursor tolerance, and the fragment ions were restricted to a 600.0 ppm tolerance. All searches were parallelized and searched on the VJC proteomics cluster. Search space included all fully tryptic peptide candidates with no missed cleavage restrictions. Carbamidomethylation (+57.02146) of cysteine was considered a static modification. We required 1 peptide per protein and both tryptic termini for each protein identification. The ProLuCID search results were assembled and filtered using the DTASelect program (Cociorva et al., 2007; Tabb et al., 2002) with a peptide false discovery rate (FDR) of 0.001 for single peptides and a peptide FDR of 0.005 for additional peptides for the same protein. Under such filtering conditions, the estimated false discovery rate for peptides was never more than 0.5%. Spectra for individual posttranslational modifications of interest were manually inspected.

### Online supplemental material

Movie S1 shows the Htb1-mCherry marker strain undergoing meiosis, Movie S2 the GFP-Spo20^51-91^ marker strain, and Movie S3 the Spc42-GFP strain. Movie S4 and Movie S5 show *pGAL-NDT80* (± *GAL* induction) cells undergoing meiosis or arrested in prophase I. Movie S6 and Movie S7 show *ime2-as1* and control cells undergoing meiosis. Figure S1 contains montages of Movies S4-S7. Figure S2 shows that *CDC20* is not required for Num1 degradation and that the *rpn6-1* allele phenocopies MG-132 treatment. Figure S3 shows that the *MDM36-3V5* allele is functional. Table S1 lists the strains used in this study. Table S2 lists plasmids used.

## ACKNOWLEDGMENTS

We thank Gloria Brar, Jingxun Chen, Leon Chan, David Drubin, Devon Harris, James Olzmann, Michael Rape, Tina Sing, Amy Tresenrider, and all members of the Ünal and Brar laboratories for experimental suggestions and comments on this manuscript. This work was supported by funds from the Pew Charitable Trusts (00027344), Damon Runyon Cancer Research Foundation (35-15), National Institutes of Health (DP2 AG055946-01), and Glenn Foundation for Medical Research to EÜ and a National Science Foundation Graduate Research Fellowship to EMS (DGE 1752814 and DGE 1106400). We also acknowledge technical support from Juliet Barker, Christiane Brune, Yuzhang Chen, Grant King, and Amy Tresenrider. We thank Laura Lackner for sharing the recombinant Mdm36 plasmid construct and technical help, Kevan Shokat for kindly providing the kinase inhibitors, 1-NA-PP1 and 1-NM-PP1, and Jeremy Thorner for stimulating discussions.

The authors declare no competing financial interests.

Author contributions: EÜ and EMS designed research. EMS performed experiments. EMS, PRJ, and EÜ analyzed data. LEB purified Ime2-st from yeast cells, provided technical insight, and helped with manuscript revisions. EÜ and EMS wrote the original draft and revised the manuscript.

---

**Figure S1.**
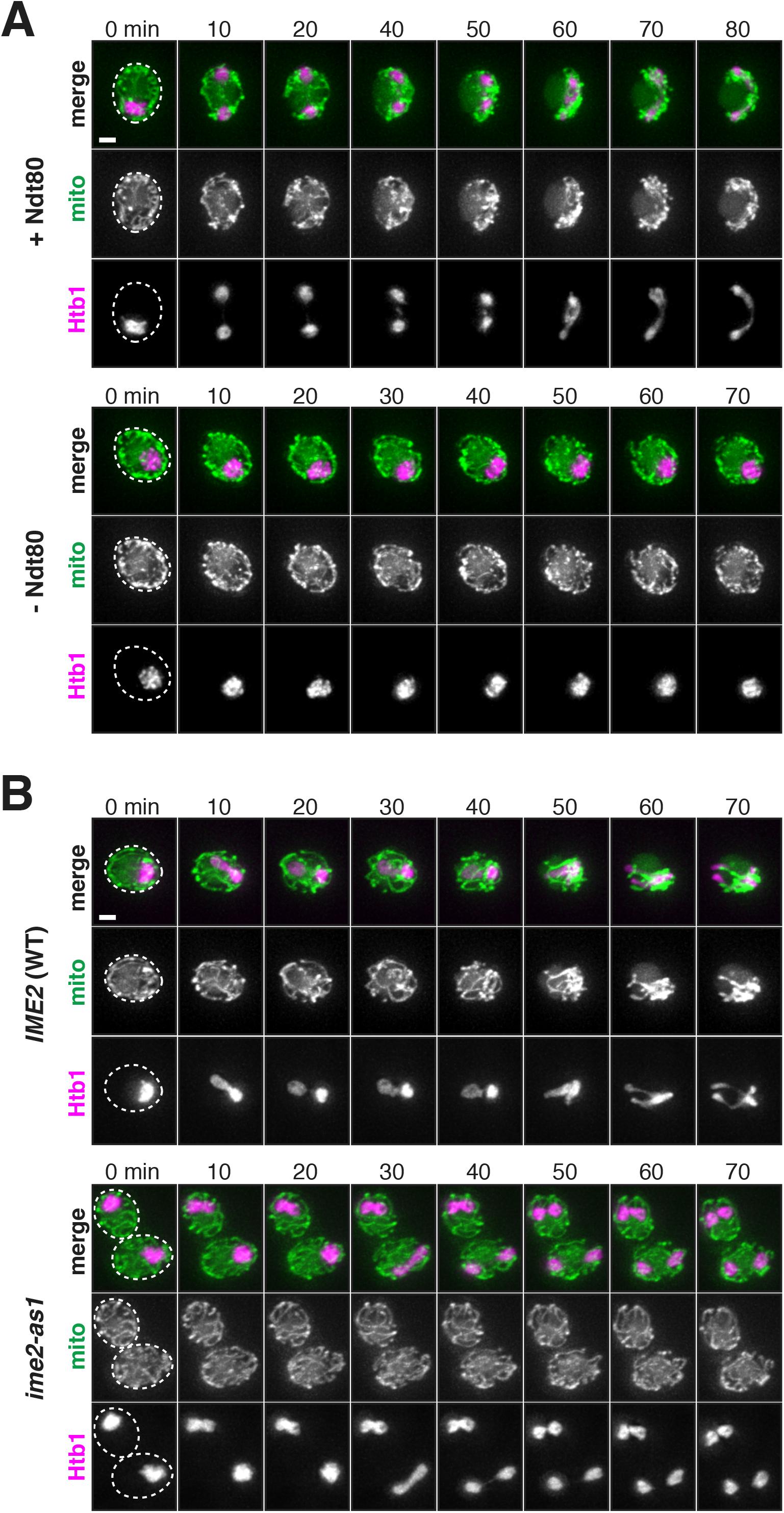
Ndt80 and Ime2 are required for mitochondrial detachment. Related to Figure 3. **A-B.** Movie montages of cells undergoing meiosis. Cells express Cit1 -GFP to label mitochondria and Htb1 -mCherry to label the nucleus. After the indicated treatments, cells were transferred to a 96-well plate and imaged every 10 min. Dashed lines indicate cell boundaries. **A.** Movie montages of *pGAL-NDT80 GAL4.ER* (UB9158) cells. After 5 h in SPO medium, the culture was split, and cells were treated with 1 *μ*M β-estradiol (+Ndt80) or ethanol vehicle control (-Ndt80). **B.** Movie montages of *pGAL-NDT80 GAL4.ER* cells carrying wild-type *IME2* (UB9158) or the *ime2-as1* allele (UB16888). After 5 h in SPO medium, cells were simultaneously treated with 1 *μ*M β-estradiol and 20 *μ*M 1-NA-PP1. Scale bar, 2 *μ*m.

**Figure S2.**
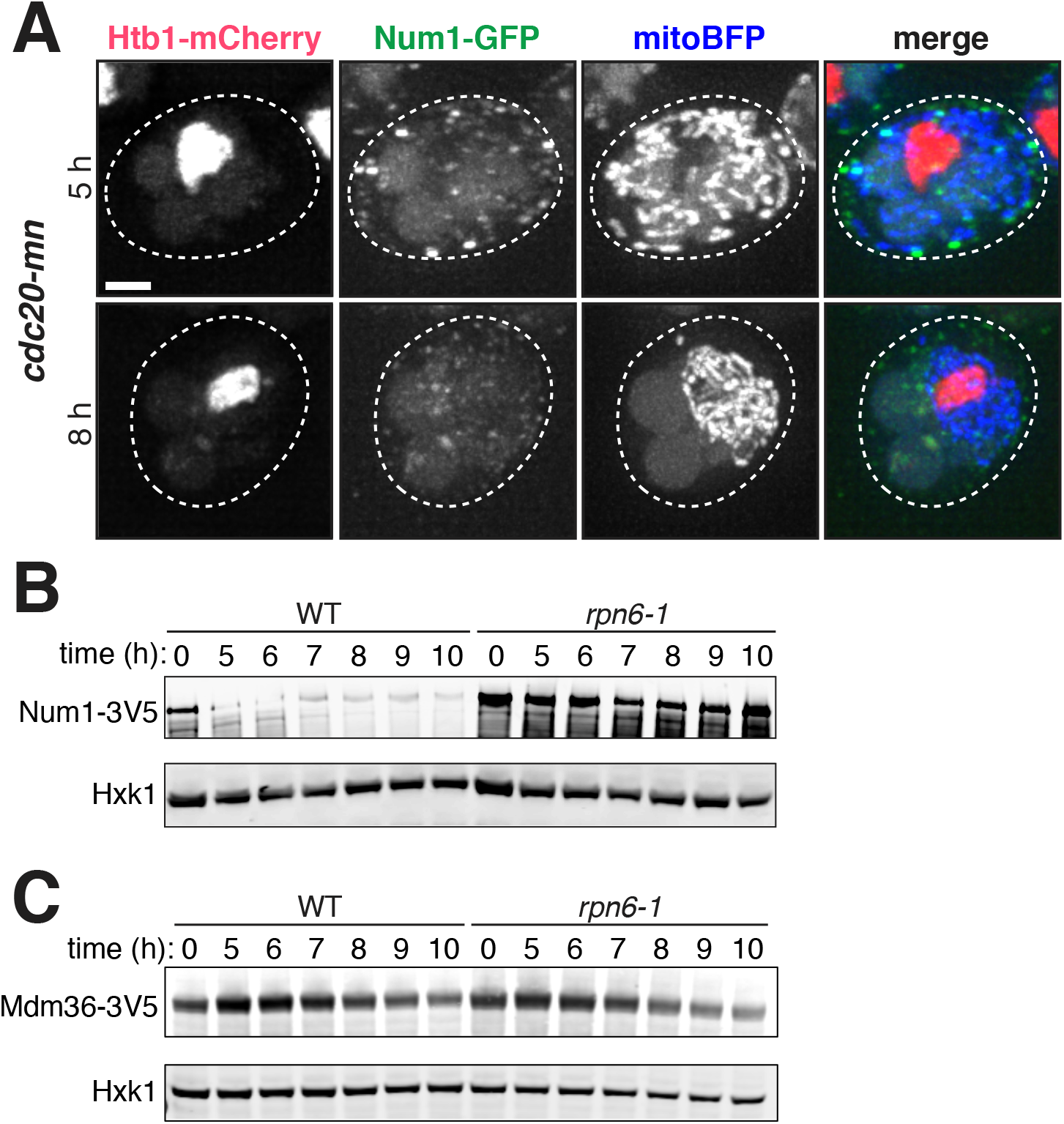
Genetic requirements for MECA subunit degradation. Related to Figures 5 and 6. **A**. Maximum intensity projections of live *pGAL-NDT80 GAL4.ER cdc20-mn* (UB16156), which is *pCLB2-CDC20,* imaged after 5 h or 8 h in sporulation media. *NDT80* expression was induced at 5 h by the addition of 1 *μ*M β-estradiol. Mitochondria, mitoBFP. Nuclei, Htb1-mCherry. **B-C.** Immunoblots of MECA subunits in *rpn6-1* mutant background and wild-type control. Strains were induced to sporulate, then 5 h later 1 *μ*M β-estradiol was added to induce *NDT80.* Hxk2 served as a loading control. **B.** Num1-3V5 wild type (UB12402) and *rpn6-1* (UB13816). **C.** Mdm36-3V5 wild type (UB13851) and *rpn6-1* (UB14340). Scale bar, 2 *μ*m.

**Figure S3.**
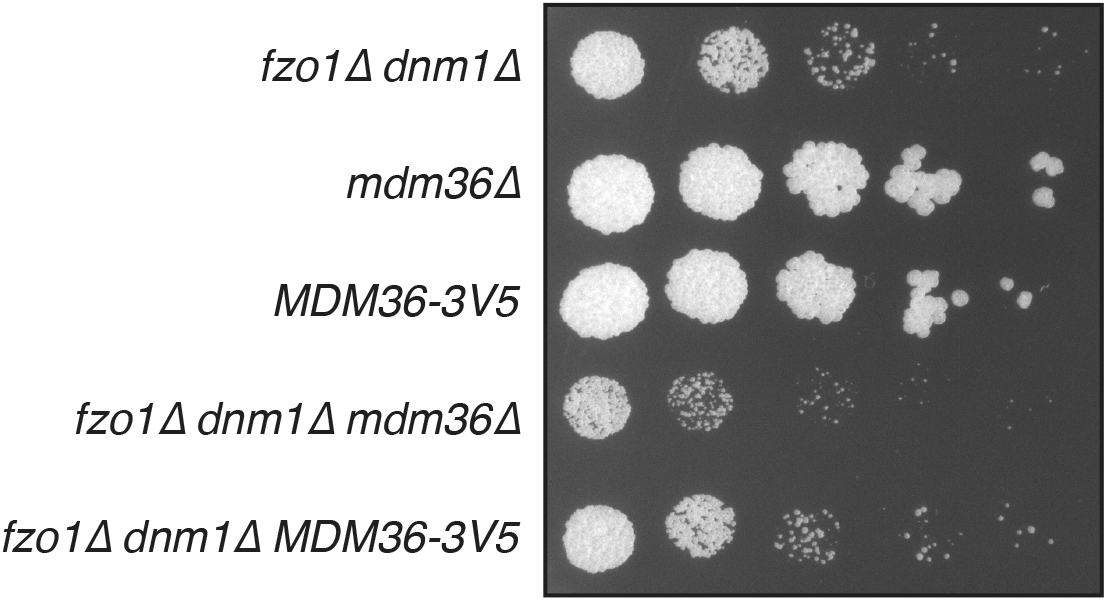
*MDM36-3V5* allele is functional. Related to Figure 5. *MDM36-3V5* does not exhibit a synthetic growth defect with the *fzo1Δ dnm1Δ* double mutant, a sensitized background to assess Mdm36 function (Lackner et al., 2013). *MDM36-3V5* (UB13762) and *mdm36Δ* (UB13770) grow normally. *fzo1Δ dnm1Δ* (UB7798) shows diminished growth on YEP + 3% glycerol, and this growth defect is more severe in *fzo1Δ dnm1Δ mdm36Δ* (UB13766), as previously described (Lackner et al., 2013). No synthetic growth defect is observed in *fzo1Δ dnm1Δ MDM36-3V5* (UB13768). Cells were pre-grown overnight on YPG plates, then serially diluted and spotted on the YPG plate shown. The plate was incubated at 30°C for 3 d.

**Table S1.**
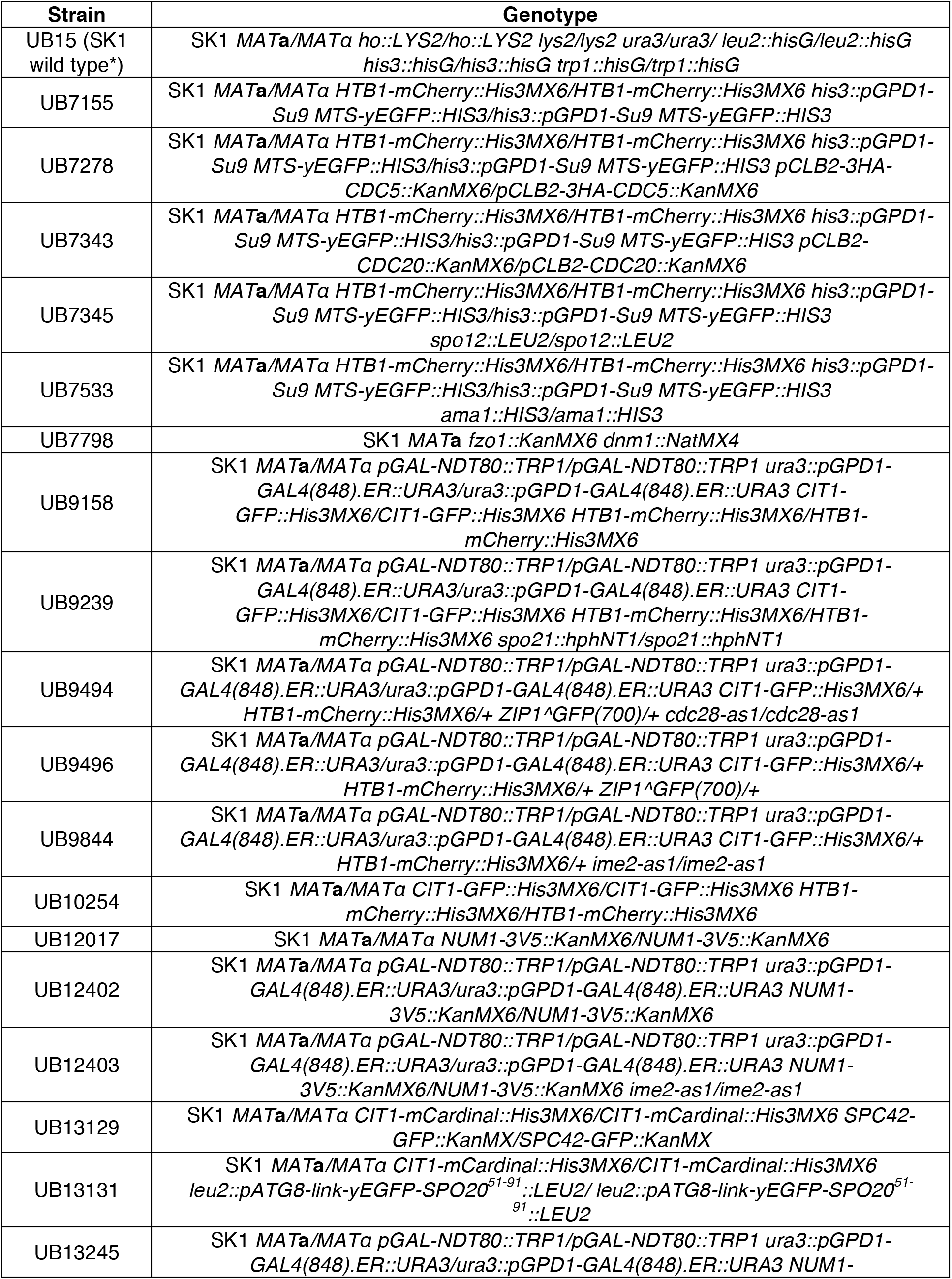
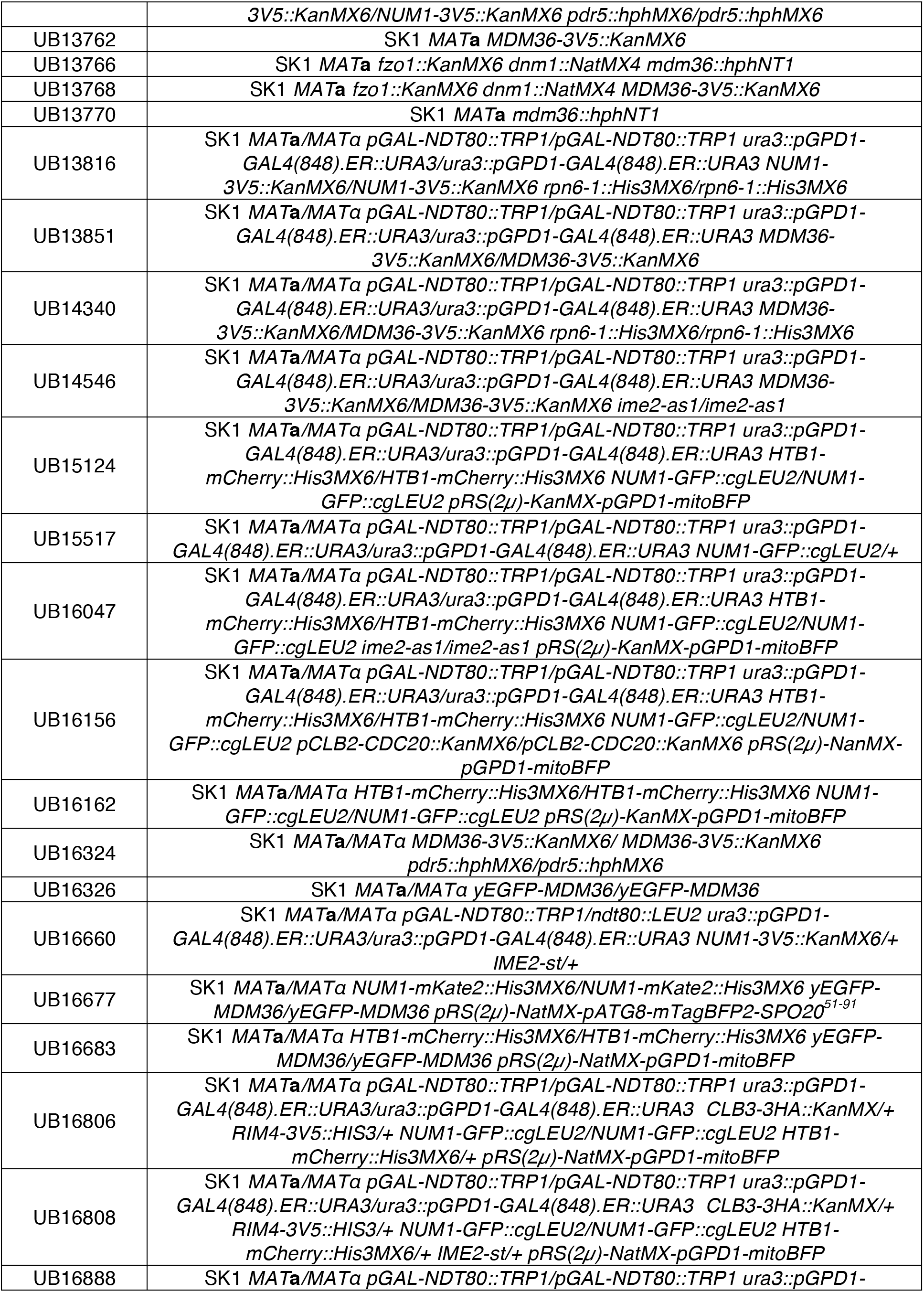
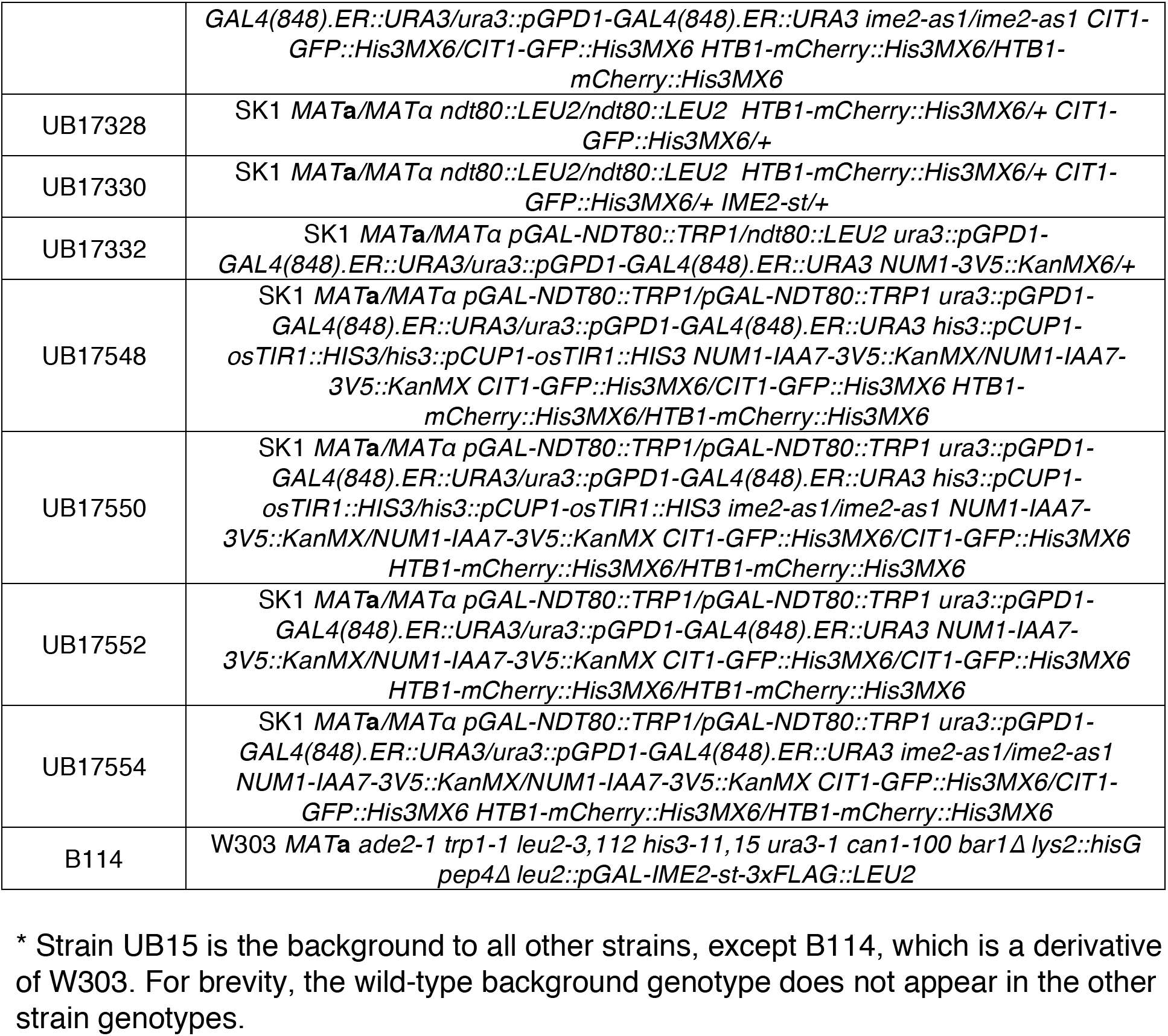
Strains used in this study.

**Table S2.** Plasmids used in this study.

| **Plasmid Name** | **Description** |
| --- | --- |
| pUB1 | pFA6a-KanMX6 |
| pUB153 | pFA6a-NatMX4 |
| pUB217 | pFA6a-hphNT1 |
| pUB294 | 3V5-KanMX6 tagging plasmid |
| pUB679 | pNH603-pGPD1-Su9 MTS-yEGFP |
| pUB763 | pFA6a-IAA7-3V5-KanMX6 |
| pUB867 | pFA6a-GFP(S65T)-His3MX6 |
| pUB868 | pFA6a-mCardinal-His3MX6 |
| pUB949 | pLC605-pATG8-link-yEGFP-SPO20<sup>51-91</sup> |
| pUB972 | pRS(2μ)-KanMX-pGPD1-mitoBFP |
| pUB976 | pNH603-pCUP1-OsTIR1 |
| pUB1004 | pFA6a-GFP(S65T)-cgLEU2 |
| pUB1375 | pRS(2μ)-NatMX-pGPD1-mitoBFP |
| pUB1392 | pRS(2μ)-NatMX-pATG8-mTagBFP2-SPO20<sup>51-91</sup> |
| pUB1395 | URA/CEN-Cas9-MDM36(gRNA3) |
| pUB1429 | pET22b mod T7prom::H6-T7-Mdm36 |

**Movie S1.** Related to Figure 1A. Movie showing the cell in Figure 1A undergoing meiosis. Htb1-mCherry (green) labels nuclei, and Cit1-GFP (magenta) labels mitochondria. The movie plays at 4 frames per second, with frames acquired every 10 min. Maximum intensity projections are shown.

**Movie S2.** Related to Figure 1B. Movie showing the cell in Figure 1B undergoing meiosis. GFP-Spo20^51-91^ (green) labels the prospore membrane (and initially, the plasma membrane), and Cit1-mCardinal (magenta) labels mitochondria. The movie plays at 4 frames per second, with frames acquired every 10 min. Maximum intensity projections are shown.

**Movie S3.** Related to Figure 1C. Movie showing the cell in Figure 1C undergoing meiosis. Spc42-GFP (green) labels spindle pole bodies, and Cit1-mCardinal (magenta) labels mitochondria. The movie plays at 4 frames per second, with frames acquired every 10 min. Maximum intensity projections are shown.

**Movie S4.** Related to Figure S1A. Movie showing the cell in Figure S1A (+Ndt80) undergoing meiosis. Htb1-mCherry (magenta) labels nuclei, and Cit1-GFP (green) labels mitochondria. The movie plays at 4 frames per second, with frames acquired every 10 min. Maximum intensity projections are shown.

**Movie S5.** Related to Figure S1A. Movie showing the cell in Figure S1A (-Ndt80) undergoing meiosis. Htb1-mCherry (magenta) labels nuclei, and Cit1-GFP (green) labels mitochondria. The movie plays at 4 frames per second, with frames acquired every 10 min. Maximum intensity projections are shown.

**Movie S6.** Related to Figure S1B. Movie showing the cell in Figure S1B (*IME2* WT) undergoing meiosis. Htb1-mCherry (magenta) labels nuclei, and Cit1-GFP (green) labels mitochondria. The movie plays at 4 frames per second, with frames acquired every 10 min. Maximum intensity projections are shown.

**Movie S7.** Related to Figure S1B. Movie showing the cell in Figure S1B (*ime2-as1*) undergoing meiosis. Htb1-mCherry (magenta) labels nuclei, and Cit1-GFP (green) labels mitochondria. The movie plays at 4 frames per second, with frames acquired every 10 min. Maximum intensity projections are shown.

